# The state of oligomerization of Rubisco controls the rate of LSU translation in *Chlamydomonas reinhardtii*

**DOI:** 10.1101/2020.10.21.348813

**Authors:** Wojciech Wietrzynski, Eleonora Traverso, Francis-André Wollman, Katia Wostrikoff

## Abstract

Ribulose 1,5-bisphosphate Carboxylase/Oxygenase (Rubisco) is a key enzyme for photosynthesis-driven life on Earth. While present in all photosynthetic organisms, its most prominent form is a hetero-oligomer in which a Small Subunit (SSU) stabilizes the core of the enzyme built from Large Subunits (LSU), yielding, after a chaperone-assisted multistep assembly, a LSU_8_SSU_8_ hexadecameric holoenzyme. Here we use *Chlamydomonas reinhardtii*, and a combination of site-directed mutants, to dissect the multistep biogenesis pathway of Rubisco *in vivo*. We identify assembly intermediates, in two of which LSU is associated with the RAF1 chaperone. Using genetic and biochemical approaches we further unravel a major regulation process during Rubisco biogenesis which places translation of its large subunit under the control of its ability to assemble with the small subunit, by a mechanism of Control by Epistasy of Synthesis (CES). Altogether this leads us to propose a model where the last assembly intermediate, an octameric LSU_8_-RAF1 complex which delivers LSU to SSU to form the Rubisco enzyme, converts to a key regulator form able to exert a negative feed-back on the initiation of translation of LSU, when SSU is not available.

## Introduction

Ribulose bisphosphate carboxylase/oxygenase (Rubisco) is the key enzyme in the light-driven carbon assimilation pathway. Emerging around 3.5 billion years ago, even before the beginning of oxygen-evolving photosynthesis, it is one of the most abundant proteins on Earth (Ellis, 1979; Tabita et al., 2008; Bar-On and Milo, 2019). Operating as the first step in the Calvin-Benson-Basham cycle, Rubisco catalyzes the fixation of atmospheric CO_2_ into, biologically available, organic carbon. Throughout time, Rubisco evolved in different forms which are now present in all photosynthetic organisms (Andersson and Backlund, 2008; Tabita et al., 2008). The most widespread clade, form I, consists of Rubisco formed by both a Large (LSU) (~52 kDa) and a Small (SSU) (~16 kDa) subunits. Rubisco form IB, a further subclade division, is present in cyanobacteria as well as in green algae and plants (Hauser, 2015). In the latter eukaryotic organisms the two subunits are encoded by spatially separated genomes of different ploidy. LSU is encoded by a single gene (*rbcL*) in the highly polyploid chloroplast, whereas SSU is a product of a family of nuclear genes (*RBCS*). Both subunits assemble in a 1:1 ratio in the chloroplast stroma, to create a hexadecameric holoenzyme (Andersson and Backlund, 2008).

Recently, significant progress was made in the comprehension of the mechanisms leading to Rubisco biogenesis in the chloroplast (reviewed in (Bracher et al., 2017; Vitlin Gruber and Feiz, 2018)). They rely on conserved features observed for cyanobacterial Rubisco assembly, while exhibiting eukaryotic specificities. As mentioned above, a first eukaryotic feature of green algae and vascular plants consists in the transfer of *RBCS* genes to the nucleus, no longer clustered with the *rbcL* gene in an operon. This allows for further regulatory processes to take place. *RBCS* genes were indeed early characterized as being part of the PhANG gene family (reviewed in (Chan et al., 2016; Börner, 2017)), a set of genes undergoing retrograde signaling in response to chloroplast translation and redox status. Once translated in the cytosol, SSU is translocated into the chloroplast via the Tic/Toc import machinery (Jarvis, 2008) where it undergoes cleavage of its targeting peptide and post translational Met1 modification (Grimm et al., 1997). Then, it would interact with the RAF2 chaperone (Rubisco accumulation factor-2), an inactive form of pterin carbinolamine dehydratase, which would deliver the protein to an LSU oligomeric complex for its proper assembly in the final Rubisco holoenzyme (Feiz et al., 2014), as evidenced by RAF2 and SSU coimmunoprecipitation and by the decreased Rubisco accumulation observed in maize mutants. The role of RAF2 in SSU chaperoning is further supported by the observation of Arabidopsis mutants presenting a decrease in SSU production which also display reduced accumulation of RAF2 (Fristedt et al., 2018).

LSU biogenesis on the other hand starts with the expression of the chloroplast *rbcL* gene, which relies on a eukaryote specific factor. The MRL1 factor, a nuclear-encoded organellar transacting factor belonging to the pentatricopeptide repeat protein family (Woodson and Chory, 2008; Barkan and Small, 2014; Hammani et al., 2014), contributes to the stabilization of the *rbcL* transcript in Chlamydomonas and of its processed form in Arabidopsis (Johnson et al., 2010). Because of the hydrophobic nature of the LSU surface, making it aggregation-prone, nascent LSU proper folding requires the assistance of molecular chaperones. Based on a model derived for bacterial studies (Langer et al., 1992; Hartl, 1996), it was suggested that newly synthesized LSU is recruited by the chloroplast folding machinery, interacting first with the plastid homologs of the DnaK/DnaJ/GrpE chaperones and subsequently with the CPN60/CPN23/CPN10 chaperonin complex (Brutnell et al., 1999). However, despite the requirement for the DnaK/DnaJ/GrpE chaperones in the recombinant expression of Rubisco in bacteria (Checa and Viale, 1997), there is no *in vivo* experimental evidence that LSU expressed in the chloroplast is indeed a client protein for those factors, which may suggest that neosynthetized LSU is directly captured by the chaperonin complex. LSU would then oligomerize in a step-wise manner to create an octameric core of the enzyme with the help of other assembly chaperones. Two of these chaperones, RBCX (Onizuka et al., 2004) and RAF1 (Feiz et al., 2012), are of cyanobacterial origin where they have been shown to stabilize LSU during dimerization and further oligomerization until the binding of SSU, thereby leading to a displacement of the chaperones as demonstrated *in vitro* (Liu et al., 2010; Bracher et al., 2011; Hauser et al., 2015; Kolesinski et al., 2017). While RBCX and RAF1 were shown to promote folding and assembly of the LSU_8_ core *in vitro*, experimental evidence to support their role in LSU oligomerization *in vivo* is lacking, even though LSU can be identified as a major interactant for both chaperones. Indeed, LSU co-immunoprecipitates with Strep-tagged RBCXs from Arabidopsis extracts indicating that RBCX may bind to plant LSU as well (Kolesinski et al., 2011). Similarly, LSU is co-immunoprecipitated with RAF1 in maize extracts (Feiz et al., 2012) or captured from cyanobacterial extracts when mixed with a recombinant RAF1 Strep-tagged variant (Kolesinski et al., 2014; Kolesinski et al., 2017). Whether RAF1 and RBCX are functionally redundant is still under debate. RBCX is dispensable *in vivo*, at least in the β-cyanobacterial species in which this gene does not cluster within the Rubisco operon such as in *Synechococcus elongatus* sp. PCC7942 (Emlyn-Jones et al., 2006). Whether this dispensability still holds true for cyanobacterial species harboring a Rubisco LXS operon awaits further confirmation (Onizuka et al., 2004; Emlyn-Jones et al., 2006; Tarnawski et al., 2008). To date, there is no evidence of its requirement in algae and plants, where two RBCX isoforms, which would form homodimers, have been described (Kolesinski et al., 2013; Bracher et al., 2015). The requirement for RAF1 also varies between species: a RAF1 knockout is lethal for maize seedlings and results in Rubisco deficiency (Feiz et al., 2012). Moreover a cognate RAF1 is required for plant Rubisco assembly, indicating a tight LSU-RAF1 coevolution (Whitney et al., 2015). In sharp contrast, RAF1 absence in cyanobacteria still allows Rubisco formation, as monitored in *Synechocystis* sp. PCC 6803 (Kolesinski et al., 2017) and in *Synechococcus elongatus* PCC7942. In the latter, Rubisco amount was decreased by one third, followed by a default in carboxysome formation, resulting in reduced cell growth in air (Huang et al., 2020). Interestingly, while both RAF1 and RBCX appear as non-essential in cyanobacteria and seem to be able to interact *in vitro* with LSU oligomers of the same order, their mode of action may be different: the resolution of LSU-containing crystals (Bracher et al., 2011; Hauser et al., 2015; Huang et al., 2020; Xia et al., 2020) indicates indeed that RBCX and RAF1 have different binding sites on LSU, and may work sequentially, as RAF1 is able to displace RBCX from LSU while the converse is not true (Kolesinski et al., 2017). Secondly, the respective knock-outs observed in cyanobacteria exhibit different phenotypes regarding Rubisco amount and carboxysome formation, casting doubts about their possible redundancy (Huang et al., 2019; Huang et al., 2020).

A second eukaryote-specific factor is required for Rubisco biogenesis, namely the BSD2 (Bundle Sheath Defective 2) protein, first identified in maize (Brutnell et al., 1999). While the role of BSD2 remained for a long time hypothetical, a breakthrough study highlighted the requirement for the BSD2 chaperone for higher plant Rubisco recombinant production in order to stabilize LSU at the final stage of Rubisco formation before SSU binding (Aigner et al., 2017). In addition, *in vivo* data indicates that tobacco BSD2 comigrates with Rubisco (Conlan et al., 2019) suggesting that a BSD2-LSU complex is the end-state intermediate in plants. Additional roles for BSD2 in chloroplast coverage and/or division (Salesse et al., 2017; Li et al., 2020) have been proposed in maize, but may be restricted to C4 plant lineages, as they weren’t observed in tobacco (Conlan et al., 2019). Last BSD2 may also participate in a repair mechanism of oxidized Rubisco (Busch et al., 2020). There is little evidence for an actual ortholog of BSD2 in green algae. In Chlamydomonas, the ZNJ2 and ZNJ6 proteins share with plant BSD2, an homology restricted to a Zn finger domain characteristic of DNAJ-like proteins (Doron et al., 2014; Doron et al., 2018) but they display a twin arginine motif in their N-terminal domain which is typical of luminal targeting sequence lumen (Chaddock et al., 1995). There is, as yet, no report of their actual role in Rubisco biogenesis.

Noteworthy, the SSU-chaperone RAF2 also interacts with LSU *in vivo* (Feiz et al., 2014) and is required in the recombinant production of plant Rubisco *in E. coli*, independently of SSU (Aigner et al., 2017). Last, LSU also undergoes some post-translational modifications but their occurrence along the biogenesis pathway remains to be understood (Houtz et al., 2008).

Since the two Rubisco subunits are expressed in two different compartments, there is a need for some regulatory mechanism to coordinate their production in the stoichiometric amounts required for their functional assembly. It already has been demonstrated that other multimeric photosynthetic complexes, such as Photosystems I and II (PSI, PSII respectively), Cytochrome *b*_*6*_*f* and ATP-synthase, undergo translation regulatory processes, known as CES processes, sensing their assembly state in *C. reinhardtii* chloroplasts (Kuras and Wollman, 1994; Choquet et al., 1998; Wostrikoff et al., 2004; Minai et al., 2006; Drapier et al., 2007) as well as in higher plant chloroplasts (Levey et al., 2014; Chotewutmontri and Barkan, 2020): the rate of translation of a subset of chloroplast-encoded subunits was shown to depend on the presence of their assembly partners. Accordingly, earlier observations of Chlamydomonas *RBCS* knockout mutants showed that LSU synthesis is strongly decreased (Khrebtukova and Spreitzer, 1996). Similarly, tobacco *RBCS* knock-down lines (Rodermel et al., 1996) displayed a down-regulation of LSU synthesis. It was subsequently demonstrated that unassembled LSU exerts a negative feed-back on its own translation in maize and tobacco chloroplasts (Wostrikoff and Stern, 2007; Wostrikoff et al., 2012). However, the nature of the LSU assembly intermediate which controls LSU translation has not been determined. That the CES behavior of LSU has been documented both in vascular plants and green algae makes it a case study for the evolution of the underlying regulatory mechanism.

As a first step towards this goal, we undertook a detailed molecular characterization of the LSU assembly intermediates in Rubisco biogenesis in the genetically tractable microalgae - *Chlamydomonas reinhardtii*. Using different mutants affected in Rubisco LSU and SSU subunits and impairing Rubisco assembly we were able to highlight the *in vivo* intermediates of Rubisco formation and LSU-chaperone(s) complexes. We then dissected the CES mechanism governing LSU translation in the absence of SSU and provide evidence that the CES control of *rbcL* translation relies on the specific accumulation of a hetero-oligomer, comprising at least LSU and RAF1, in absence of SSU.

## Results

### Downregulation of LSU synthesis in a Chlamydomonas RBCS mutant

Previous work showed that deletion of the *RBCS* genes in *Chlamydomonas reinhardtii* resulted in a significant decrease in synthesis of LSU (Khrebtukova and Spreitzer, 1996). We confirmed this observation in an independent mutant, hereafter called ΔRBCS strain which, in contrast to the previously described one, is fertile, thus allowing further genetic analysis. We isolated this strain from backcrosses between our laboratory reference strain and the Cal.005.013 strain (Dent et al., 2005), which harbors a large deletion encompassing the two *RBCS* linked genes. In absence of SSU, LSU accumulated to ~1% of wild type level (WT), which argues for a concerted accumulation of Rubisco subunits (Fig. 1A). Moreover, as reported by Khrebtukova and Spreitzer (Khrebtukova and Spreitzer, 1996), LSU exhibited a lower rate of synthesis in ΔRBCS compared to WT as shown by ^14^C pulse-labeling experiments (Fig. 1B). This down-regulation is posttranscriptional, as *rbcL* mRNA level was not affected in the ΔRBCS strain as compared to WT (Fig. 1C). To confirm that the decrease in LSU radiolabeling truly represents a decrease in translation rather than a massive proteolytic degradation, we tested the stability of unassembled LSU in the ΔRBCS strain by following LSU accumulation over 4 hours in presence of chloramphenicol, an inhibitor of chloroplast protein synthesis. As shown in Fig. 1D, unassembled LSU was found to be stable over this time period (see also Sup. Fig. 1B).

**Fig. 1:**
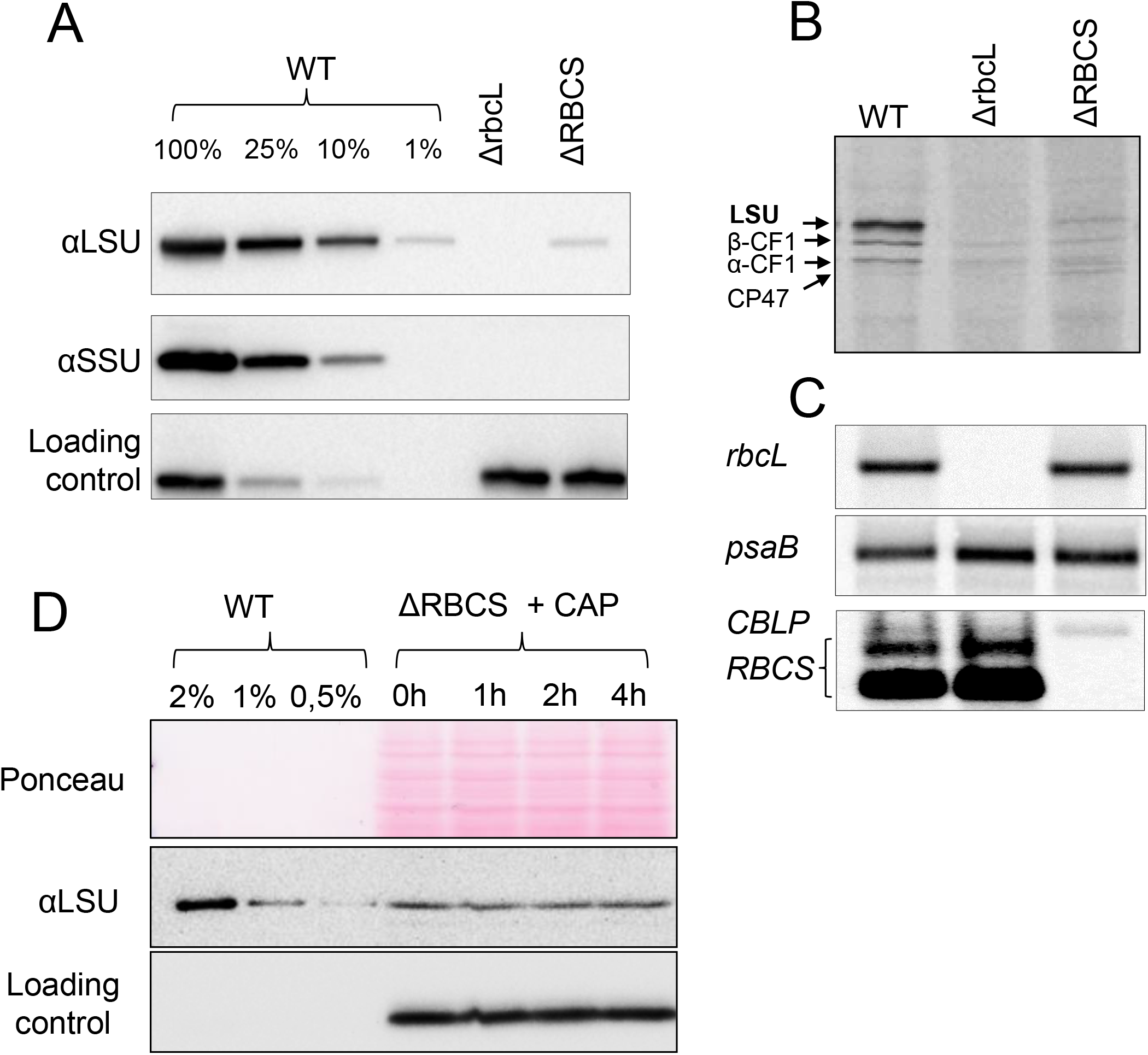
LSU accumulation, synthesis rate and stability in absence of its assembly partner. (A) Immunoblot showing protein level accumulation of Rubisco subunits in the ΔRBCS strain, using an antibody directed against whole Rubisco holoenzyme. PsaD accumulation, revealed with a specific antibody, is shown as a loading control, (B) ^14^C pulse labeling experiment showing the synthesis rate of LSU in the ΔRBCS strain as compared to the WT in upper panel (positions of LSU as well as ATPase α and β subunits and Photosystem II CP43 subunit are indicated by arrows), (C) mRNA accumulation in the same strains as in B, as probed by hybridization with *rbcL* and *RBCS* probes, and *psaB and CBLP* probes used as loading controls. In both panels, the Δ*rbcL* strain exhibiting a deletion of the *rbcL* gene is used as a negative control, (D) Unassembled LSU stability assayed by immunochase over 4 hours after chloroplast synthesis arrest by chloramphenicol (CAP) addition. LSU is detected with the anti-Rubisco antibody, cytochrome *f* is used as a loading control.

### LSU initiation of translation is impaired in absence of SSU

As shown previously, those subunits of photosynthetic complexes that undergo CES translational regulation bear, within the 5’UTR of their mRNA, all cis-acting elements controlling this process (Choquet et al., 1998; Choquet and Wollman, 2007). Thus, in all cases studied so far, regulation of translation of CES proteins occurs at the initiation step. To test whether the native *rbcL* 5’-UTR is required for the *RBCS*-sensitive down regulation of *rbcL* translation, we replaced it by the *psaA* 5’UTR. After biolistic transformation of the ΔrbcL strain (ΔR T1.3+) by the 5’UTRpsaA:*rbcL* chimera (Materials and Methods, Sup. Table 1), we obtained phototrophic transformants which accumulated WT levels of LSU, demonstrating that *psaA* 5’UTR is able to drive *rbcL* expression efficiently (Fig. 2A). We then crossed a representative 5’UTRpsaA:*rbcL* transformant (mt+) with the ΔRBCS strain (Cal.13.1B; mt-) and obtained progenies displaying uniparental inheritance of the chloroplast 5’UTRpsaA:*rbcL* chimeric gene and a 2:2 distribution of the Δ*RBCS* nuclear mutation. Progenies from distinctive genotypes (5’UTRpsaA:*rbcL* and ΔRBCS;5’UTRpsaA:*rbcL*), easily identified by their distinct acetate requirement for growth (Fig. 2A), were used in pulse labeling experiment to monitor *rbcL* translation rates in this non-native 5’UTR context. As shown in Fig. 2B, the *rbcL* translation rate was now similar between sister strains, harboring or not *RBCS*, and it proceeded at the same rate as in the WT. This experiment demonstrates that *rbcL* 5’UTR is required for CES regulation of LS synthesis since its replacement by the upstream sequence of another chloroplast gene, allows *rbcL* translation to become CES-insensitive. Yet, we note that LSU accumulation in absence of SSU is still reduced compared to the strain harbouring the 5’UTRpsaA:*rbcL* chimera and expressing SSU. However, the decrease in LSU accumulation is less pronounced than when LSU is translated from the native *rbcL* gene in absence of SSU (see discussion section).

**Fig. 2:**
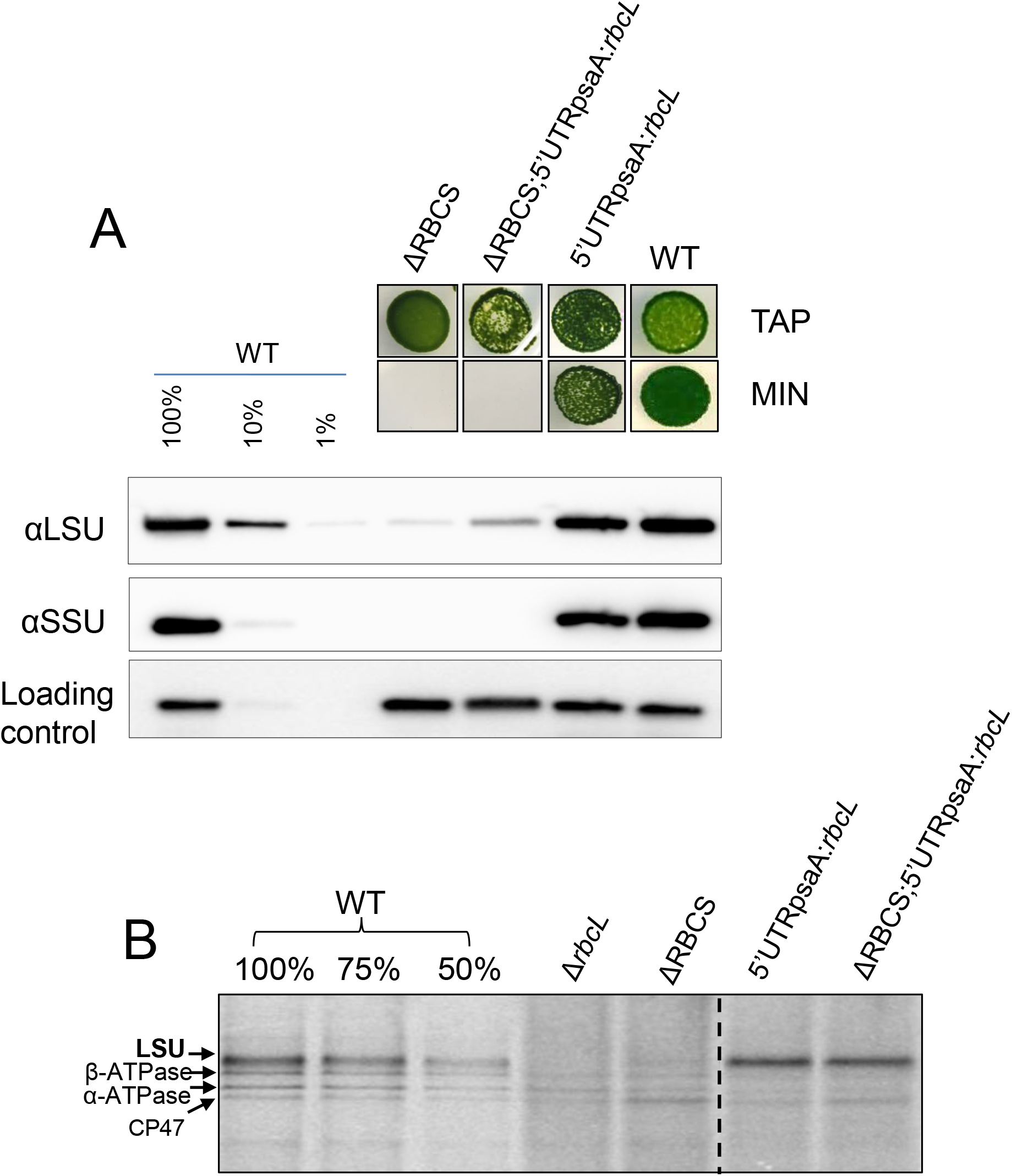
Swapping *rbcL* 5’UTR regulatory sequence impairs the CES regulation. (A) Upper panel: Photosynthetic growth phenotypes of 5’UTR*psaA*:*rbcL* strains defective or not for Rubisco SSU, and accumulation of the corresponding Rubisco subunits tested by western blot analysis. Lower panel: In ΔRBCS;5’UTRpsaA:*rbcL*, LSU is accumulating to higher levels than in ΔRBCS. TAP stands for Tris-Acetate-Phosphate medium, MIN is an acetate-free, phototrophy-selective medium. (B) ^14^C pulse labeling experiment showing *rbcL* translation rate in 5’UTRpsaA:*rbcL*-background with and without small subunit compared with wild-type, Δ*rbcL* and ΔRBCS strains. The dashed line marks the position where two irrelevant lanes were removed.

To test whether *rbcL* 5’UTR is sufficient to confer the CES regulation to an unrelated gene, we expressed a fusion between the *rbcL* 5’ UTR and the *petA* gene encoding cytochrome *f*, a core protein of the Cytochrome *b*_6_*f* complex (5’UTRrbcL:*petA*; hereafter referred to as the reporter) in presence or absence of SSU (see Materials and Methods and Sup. Table 1). Prior research (Chen et al., 1995; Choquet et al., 1998) has shown that in normal growth conditions cytochrome *f* accumulation level mirrors its rate of synthesis, which makes it a faithful reporter (proxy) of translation efficiency. In the experiment shown on Fig. 3 we compared the accumulation of the cytochrome *f* reporter protein in representative transformants in presence (5’UTRrbcL:*petA*) or absence (ΔRBCS;5’UTRrbcL:*petA*) of SSU. Expression of the reporter fusion driven by the *rbcL* 5’UTR in presence of SSU, led to significant cytochrome *f* accumulation, albeit at lower levels than observed in WT. However, cytochrome *f* accumulation driven by the *rbcL* 5’UTR became almost undetectable in absence of SSU, thus showing the same behavior as LSU. Altogether, these observations demonstrate that the 5’UTR of the *rbcL* gene contains all information required to confer a Rubisco assembly-dependent regulation of translation to a downstream coding sequence.

**Fig. 3:**
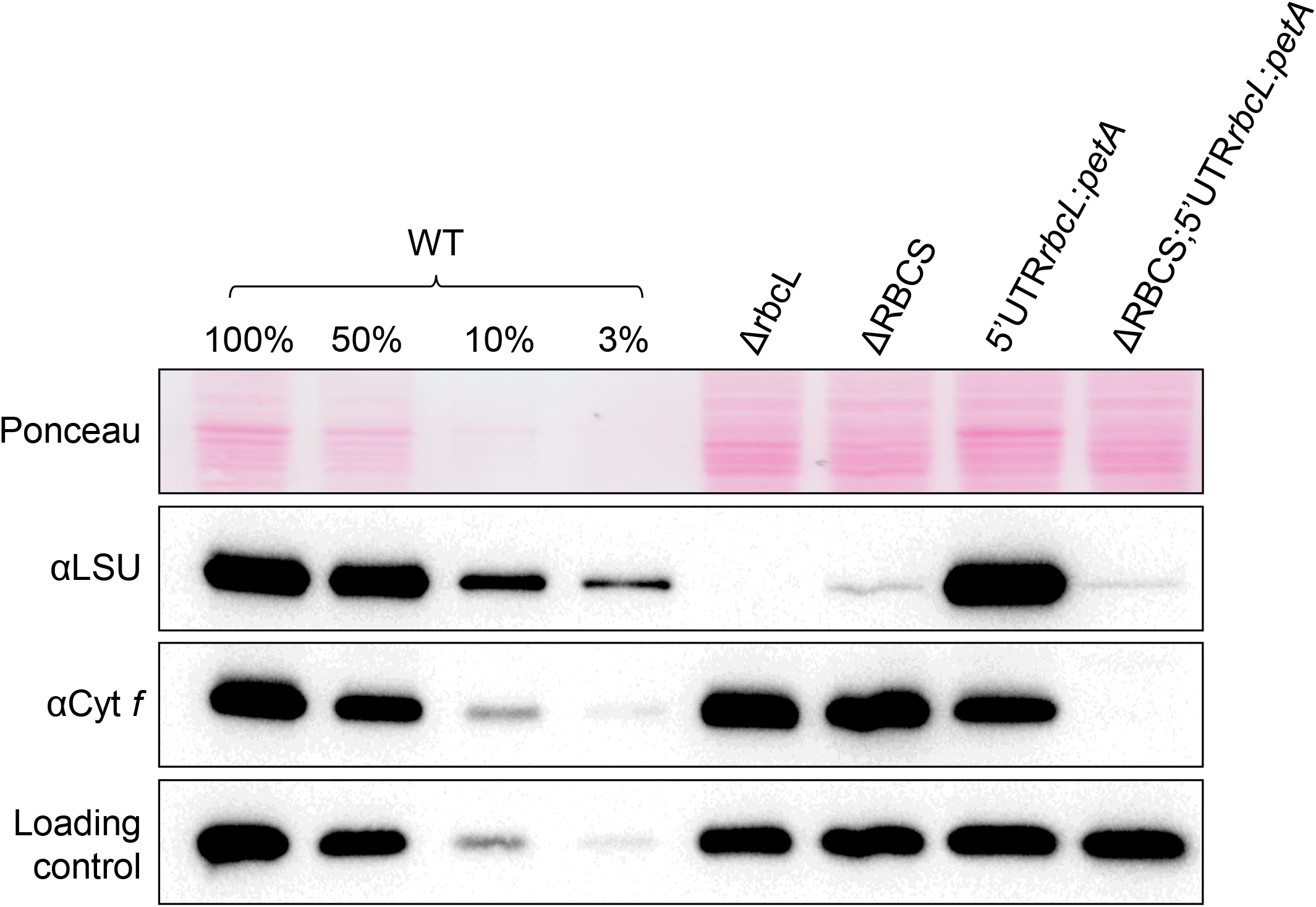
Expression of cytochrome *f* is inhibited in the absence of Rubisco small subunit. Immunoblot using antibodies directed against the proteins depicted at the left, showing Rubisco and cytochrome *f* accumulation levels in the wild-type, ΔRBCS, ΔrbcL and 5’UTRrbcL:*petA* strains with and without SSU. PsaD accumulation is shown as a loading control.

### Translation initiation is inhibited by unassembled LSU

Two possible mechanisms could account for the observed translational repression of *rbcL* in absence of SSU. Either the small subunit is required for direct or indirect trans-activation of LSU translation, or, in the absence of its SSU partner, unassembled LSU is inhibiting its own translation via an auto-regulatory feedback. In order to distinguish between the two hypotheses, we followed the expression of the *rbcL* 5’UTR-driven-reporter gene in a context where both Rubisco subunits are absent (as detailed in (Choquet and Wollman, 2007)). A trans-activation model predicts that the absence of SSU and LSU should yield a low accumulation of the reporter, similarly to what is observed in the single ΔRBCS mutant. Indeed the expression of the reporter gene should only depend on SSU in this hypothesis. Alternatively, the absence of SSU and LSU should lead to a high accumulation of the reporter in case of a negative feed-back loop. To test these two models we first generated a strain in which LSU production was prevented. To this end, a construct bearing a deletion of 116 aa was introduced at the *rbcL* locus, leading to the expression of a short, truncated polypeptide of 14 kDa composed of the N-terminal part (107 aa) fused in frame to the C-terminus (9 aa). Biolistic transformation of the ΔrbcL (ΔR T1.3+) and ΔRBCS (Cal.13.5A+) strains by the pLStr plasmid carrying this truncation (Materials and Methods and Sup. Table 1) yielded strains where truncated LSU is expressed in presence or absence of SSU (LSU_tr_ and ΔRBCS;LSU_tr_). *In vivo* pulse labeling experiments revealed that truncated LSU is robustly synthesized in the LSU_tr_ transformants (Fig. 4A). Furthermore, its synthesis rate was not altered in the absence of SSU (ΔRBCS;LSU_tr_). Yet, *rbcL* truncation led to the complete absence of LSU accumulation, which could not be detected even as trace amounts in the expected 14 kDa size range (Sup. Fig. 2). We next examined strains expressing truncated LSU in the reporter gene background with and without SSU (5’UTRrbcL:*petA* and ΔRBCS;5’UTRrbcL:*petA*) (see Materials and Methods, and Sup. Table 1). Figure 4B shows that the cyt. *f* reporter becomes expressed to a significant level in the truncated LSU background (LSU_tr;_ 5’UTRrbcL:*petA* and ΔRBCS;LSU_tr;_ 5’UTRrbcL:*petA*) compared to the strains exhibiting native LSU (5’UTRrbcL:*petA)*. Its overexpression is observed irrespective of the presence of SSU (although not quite to the same extent in the ΔRBCS;LSU_tr;_ 5’UTRrbcL:*petA* as in the LSU_tr;_ 5’UTRrbcL:*petA* strain), in sharp contrast with the ΔRBCS;5’UTRrbcL:*petA* strain in which cyt. *f* did not accumulate (Fig. 3 and 4B lane 2). Thus the second model proved correct: unassembled LSU exerts a negative feed-back on its own translation in Chlamydomonas, similar to what some of us had proposed for tobacco Rubisco (Wostrikoff and Stern, 2007).

**Fig. 4:**
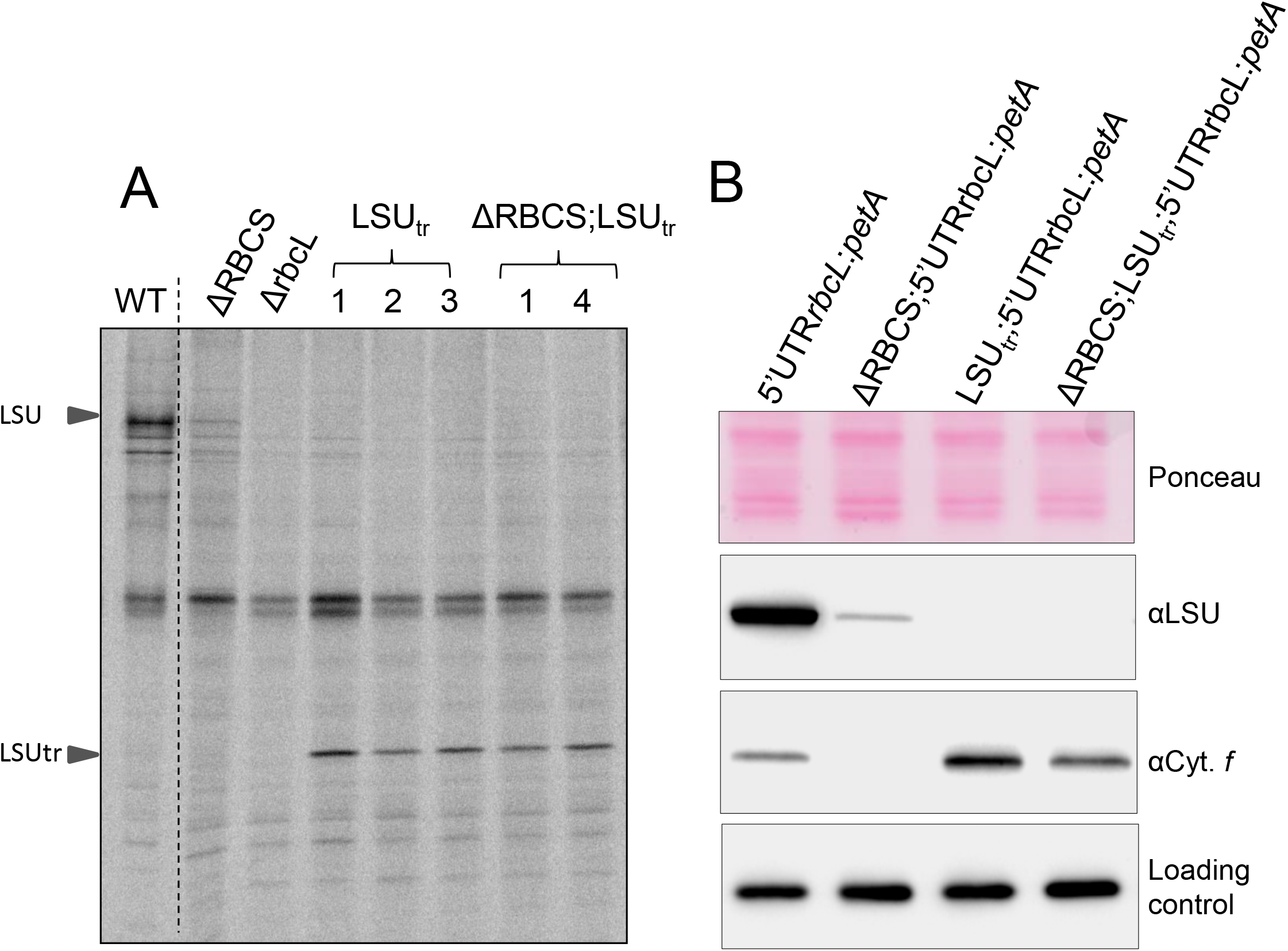
CES regulation no longer occurs in the absence of LSU accumulation. (A) ^14^C labeling experiment showing synthesis rates of chloroplast proteins in WT, ΔrbcL, ΔRBCS, LSU_tr_ transformants (1-3) and ΔRBCS;LSU_tr_ (1 and 4) strains. Migration of full-length and truncated LSU is indicated on the left. The dashed line marks the position where two irrelevant lanes were removed. (B) Immunoblot depicting LSU and cyt. *f* accumulation in representative transformants carrying both the 5’UTRrbcL;*petA* reporter gene and a truncation within the *rbcL* gene, associated or not to the ΔRBCS mutation, in comparison to the wild-type and ΔRBCS;5’UTRrbcL:*petA* strains. Ponceau stain and PsaD accumulation are shown as loading controls.

### RAF1-LSU intermediates accumulate in absence of SSU

The Rubisco assembly pathway is suggested to comprise several LSU oligomerization steps followed by SSU binding (Liu et al., 2010; Hauser et al., 2015). To investigate in which oligomerization state the unassembled, repressor-competent form of LSU accumulates, we performed a Native PAGE analysis of soluble extracts from whole cells. Immunoblotting against LSU readily detects the native Rubisco holoenzyme in a WT extract (2% dilution) (Fig. 5A). No other assembly intermediates were detected even after prolonged membrane exposure (data not shown) consistent with the idea that Rubisco assembly is a fast and dynamic process, which does not allow significant steady state accumulation of any assembly intermediate. Immunoblotting of ΔrbcL extracts revealed that the anti-Rubisco antibody cross-reacts with two LSU-unrelated bands, marked by asterisks on figure 5. These two bands are also found in the ΔRBCS extracts indicating further that they are not related to SSU either. In the absence of SSU, the residual unassembled LSU (corresponding to about ~1% of WT level, Fig. 1) partitions into three distinct LSU-reactive complexes (Fig. 5A, ΔRBCS lane). Using 2D electrophoresis and immunoblotting (Fig. 5B), we identified a band migrating above 720 kDa (depicted as a square in Fig. 5), which we attribute to the CPN60 chaperonin-bound LSU, as reported in previous studies with pea (Roy et al., 1982) and maize extracts (Feiz et al., 2012). A similar observation was reported regarding the CPN60 bacterial homolog GroEL in *in vitro* reconstitution experiments (Liu et al., 2010; Hauser et al., 2015). This attribution was confirmed with the use of a CPN60α/β1 directed antibody, which revealed two reactive bands. The upper one is found to co-migrate with the 720 kDa LSU complex, whereas the lower one migrates in the molecular mass position of free CPN60 monomers. We then suspected that the two other LSU-associated complexes observed in Figure 5A would be LSU oligomers bound to other assembly chaperones.

**Fig. 5:**
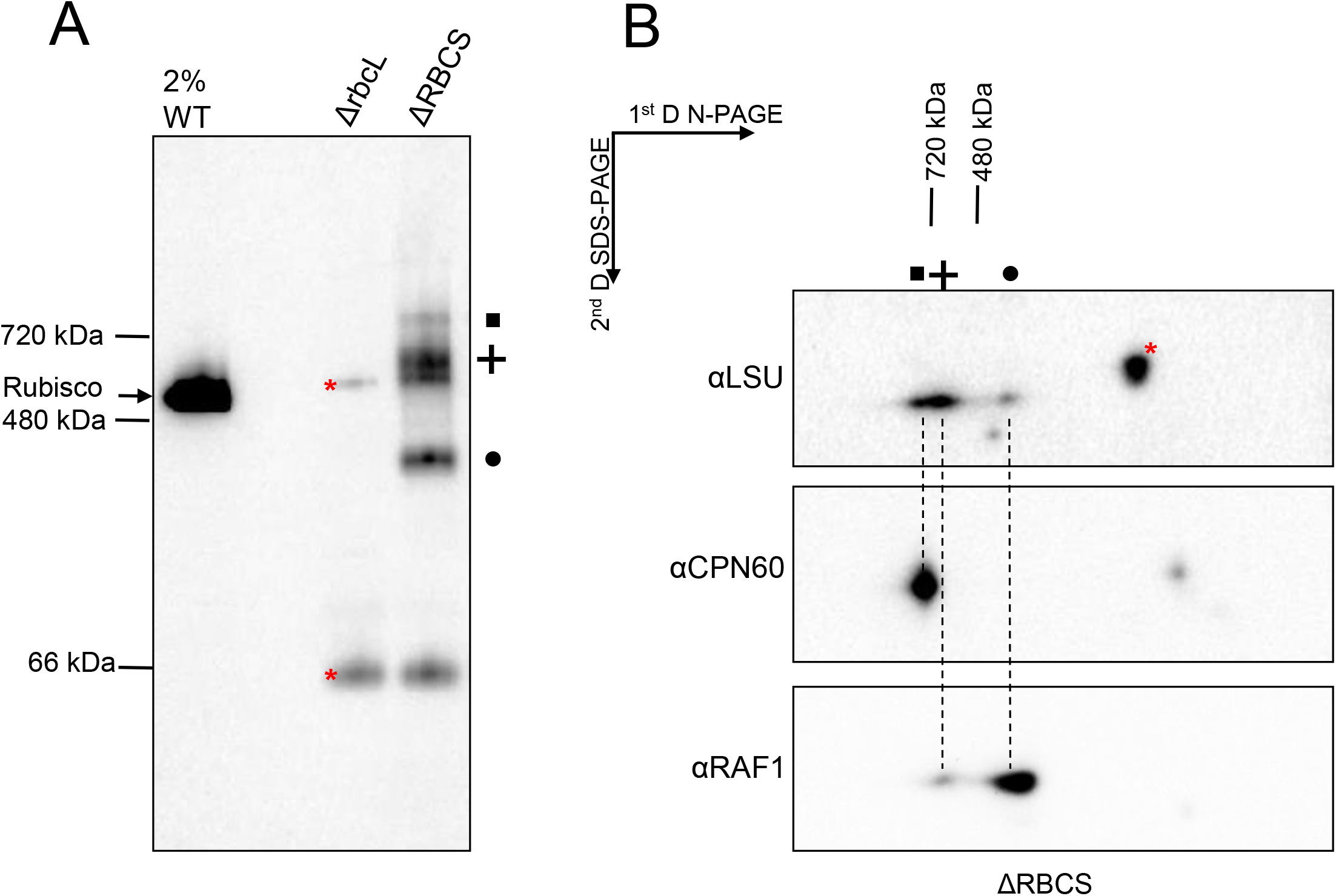
LSU assembly intermediates accumulate in the SSU-lacking strain. (A) Immunoblot with the antibody directed against LSU after Native PAGE analysis of soluble protein extracts from WT (diluted to 2% as not to obscure the gel), ΔrbcL and ΔRBCS strains. The migration of native molecular weight markers is indicated on the left. The position of Rubisco holoenzyme, as deduced from the WT signal, is indicated as well. Three LSU-specific complexes are observed in the SSU-lacking strain (depicted by a square, cross and circle). (B) Analysis of the second dimension on SDS PAGE gel by immunodetection of proteins putatively associated to LSU complexes in ΔRBCS strain (depicted by a square, cross and circle as in A), using anti-LSU, anti-CPN60 and anti-RAF1 antibodies. Dashed lines are drawn to help with the alignment. Red asterisks mark cross-contaminating signals of the anti-LSU antibody.

RBCX and RAF1 were shown to allow cyanobacterial LSU oligomerization in reconstitution experiments. We then focused on RAF1 since it has been shown to interact *in vivo* with plant LSU (Feiz et al., 2012), and is conserved in Chlamydomonas, together with RBCX (Liu et al., 2010; Bracher et al., 2011; Hauser et al., 2015). We raised an antibody directed against Chlamydomonas RAF1 (Cre06.g308450, Sup. Fig. 3) and tested whether RAF1 is able to interact with LSU in Chlamydomonas. We constructed an epitope-tagged version of RAF1 driven by a strong promoter (pJHL-RAF1S plasmid, described in Sup. Material and Methods) and transformed the ΔRBCS recipient strain. We recovered ΔRBCS;*RAF1:Strep-TG* transformants showing a three-fold overexpression of RAF1. An immunoprecipitation experiment from soluble extracts of one of these transformants and the ΔRBCS untagged control strain as then performed. After incubation with magnetic StrepTactin-coated agarose beads, the Strep-tagged RAF1 was able to efficiently pull down both tagged and untagged RAF1, as well as a significant part of LSU (Fig. 6). Thus the native and Strep-tagged RAF1 can oligomerize, either directly and/or through LSU. Altogether these experiments show that Chlamydomonas RAF1 does genuinely interact with LSU in Chlamydomonas, and that this interaction is independent of the presence of SSU, in agreement with reconstitution experiments performed with the recombinant cyanobacterial proteins (Hauser et al., 2015).

**Fig. 6:**
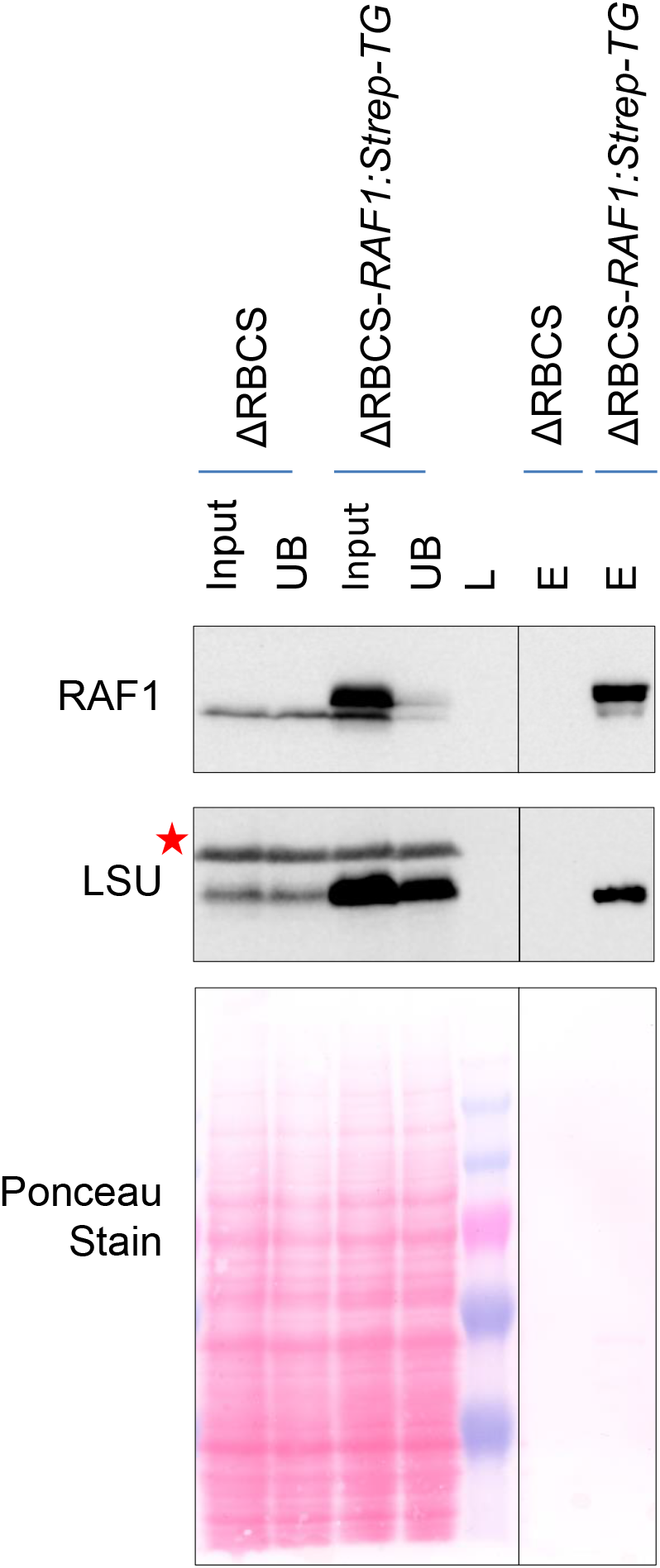
RAF1 and LSU interact in Chlamydomonas. Immunoblots showing RAF1 and LSU co-immunoprecipitation. A similar fraction of the input, unbound (UB), or bead-extracted (E) fraction from the immunoprecipitation of soluble extracts from either the ΔRBCS or ΔRBCS-*RAF1:Strep-TG* strains was separated on SDS-PAGE gel, together with a molecular weight ladder (L). The line separates non-contiguous lanes of the same gel with the same exposure. RAF1 and LSU were detected by immunoblots using specific antibodies. The anti-RAF1 antibody recognizes both the endogenous (lower band) and the overexpressed epitope-tagged RAF1 (upper band), which could be separated by this gel system. The anti-Rubisco antibody recognizes LSU as wells as an unrelated cross-reacting band marked by a red asterisk. LSU is specifically pulled-down by coimmunoprecipitation of the strep-tagged RAF1 protein. We note that not all LSU is pulled down, which could reveal the LSU fraction not associated to RAF1.

We next assessed the oligomeric state of RAF1-LSU complexes *in vivo*. After two dimensional electrophoresis and immunoblotting of soluble ΔRBCS extracts (Fig. 5B), we were able to detect a RAF1 signal co-migrating with LSU in the two complexes below the 720 kDa and 480 kDa markers (depicted respectively by a circle and a cross). We noted that most of the signal is found in the lower molecular LSU complex (hereafter called LMW-LSU), whose apparent molecular mass would be consistent with an interaction between a RAF1 dimer and an LSU dimer. Consistent with the similar accumulation of RAF1 in the WT, ΔRBCS or ΔrbcL strains (Sup. Fig. 4), both the HMW- and LMW-LSU associated RAF1 signals present in the ΔRBCS extract disappeared in the ΔrbcL extract while a new RAF1 signal was observed above the LMW-LSU position (Fig. 7). These observations argue for a genuine interaction of RAF1 with LSU in these two bands, instead of a mere co-migration. Notably we do not observe LSU-free RAF1 in absence of SSU (figure 5B and 7), neither as a monomer nor as dimer, the latter being the form observed in solution when recombinant cyanobacterial RAF1 is produced (Hauser et al., 2015; Xia et al., 2020). This, together with the immunoprecipitation data, indicates that RAF1 plays a role in LSU stabilization in Chlamydomonas as has been reported for cyanobacterial LSU in *in vitro* studies. RAF1 most probably interacts with the LSU dimer, and part of it remains associated with higher LSU oligomers *in vivo*. Interestingly, while the LSU-RAF1 dimer also accumulates in a WT background, the LSU-RAF1 HMW observed in ΔRBCS is no longer detected. Instead another RAF1 HMW oligomer, showing a different migration pattern is observed, as indicated by a star in Fig. 7. How this might relate to the Rubisco assembly process and CES autoregulation will be further discussed in the discussion section.

**Fig. 7:**
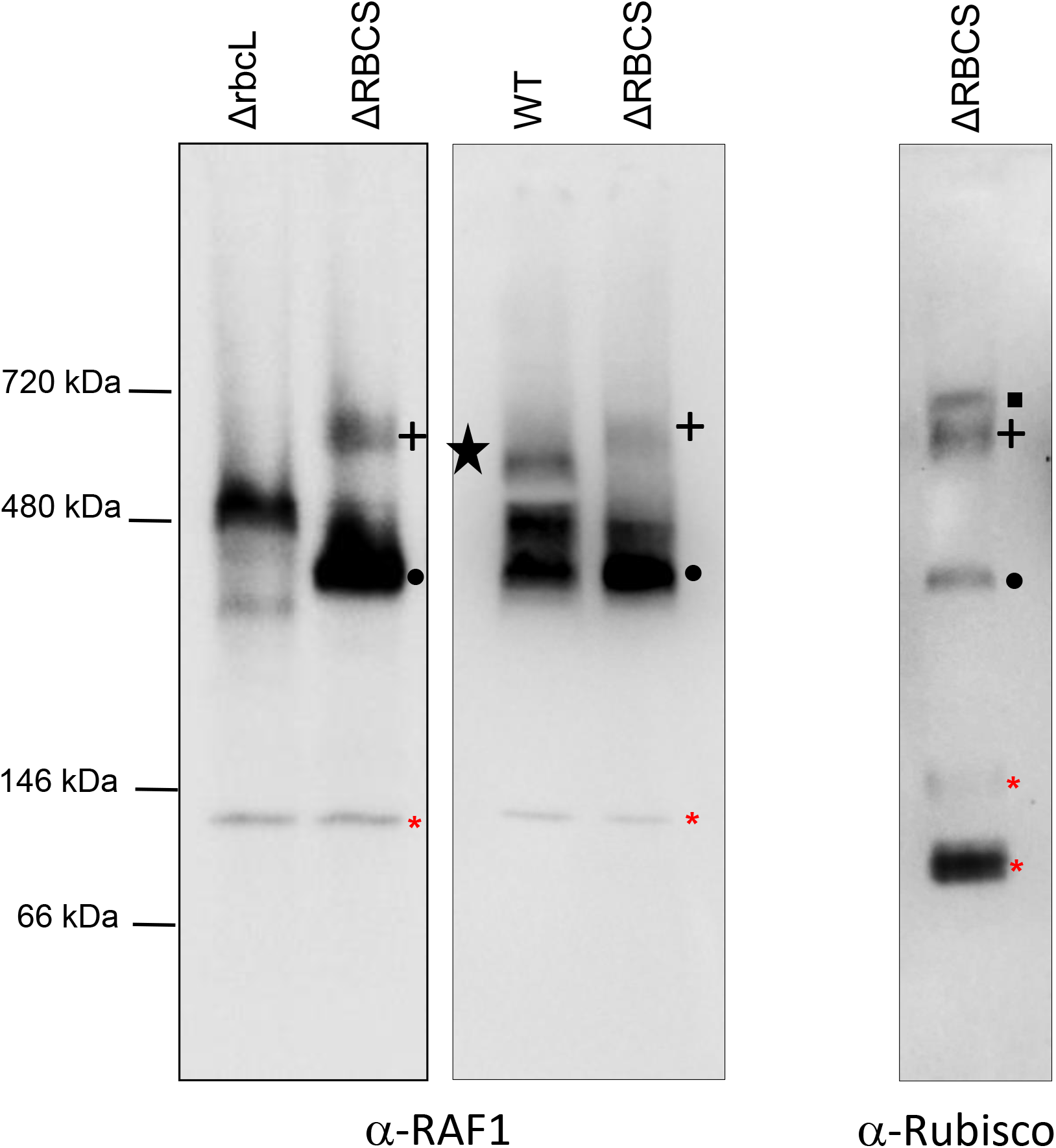
RAF1 oligomerization state in Rubisco mutants versus WT. Immunoblot of a 1D Native-PAGE of soluble extracts from ΔrbcL, ΔRBCS and WT using RAF1 (left and middle panels) or Rubisco antibody (right panel), showing that RAF1 accumulates as an oligomer in the absence of LSU. RAF1-LSU complexes are indicated using the same symbols as in Fig. 5. Note that the RAF1-LSU HMW complex found in the ΔRBCS is no longer detected in a WT background, whereas an additional low abundant RAF1 complex, indicated by a black star, is found. Red asterisks indicate antibody cross-reacting bands. (The left panel is a distinct experiment from the middle and right panels, who were separated on the same gel).

### CES regulation no longer occurs in Rubisco oligomerization mutants

To determine which LSU assembly intermediate is involved in the CES behavior of Rubisco, we undertook a structure-guided mutagenesis approach. LSU dimer interaction involves two stabilizing salt bridges between the E109 and R253 residues, and between the E110 and R213 residues from head-to-tail adjacent LSU monomers (Bracher et al., 2011) (see Sup. Fig. 5A). To alter the formation of LSU dimers or their stabilization, we introduced two substitutions within the *rbcL* gene (E109A and R253A), aimed at preventing the formation of one of these salt bridges linking LSU monomers. The resulting LSU_2_mut (*rbcL*_E109A-R253A_) transformants were no longer phototrophic, as shown for a representative transformant in Fig. 8A. One of these transformants (in mt+ background) was further crossed to ΔRBCS (mt-) to obtain a strain expressing these two *rbcL* substitutions in the absence of SSU. The resulting double LSU_2_mut-SSU mutant accumulates soluble LSU at comparable level to that in the single ΔRBCS mutant (Fig. 8A). To monitor the rate of LSU synthesis in the dimerization mutants we performed *in vivo* pulse-labeling experiment with ^14^C acetate as shown in Fig. 8B for two representative progenies of the cross. In both LSU_2_mut and ΔRBCS;LSU_2_mut genotypes, the LSU translation rate was higher than WT levels, which is indicative of deregulated translation even in total absence of SSU. Further characterization by Native PAGE and immunoblotting demonstrated that these mutations prevented accumulation of any LSU assembly intermediates beside the original LSU-CPN60 complex which in both genotypes is more abundant than in ΔRBCS (Fig. 8C). No monomeric LSU could be detected, suggesting that dimerization is required to generate the chaperonin-independent, stable forms of LSU observed in Fig. 5. We conclude that accumulation of some assembly intermediate, downstream of the CPN60-bound LSU complex is required for the CES regulation to occur.

**Fig. 8:**
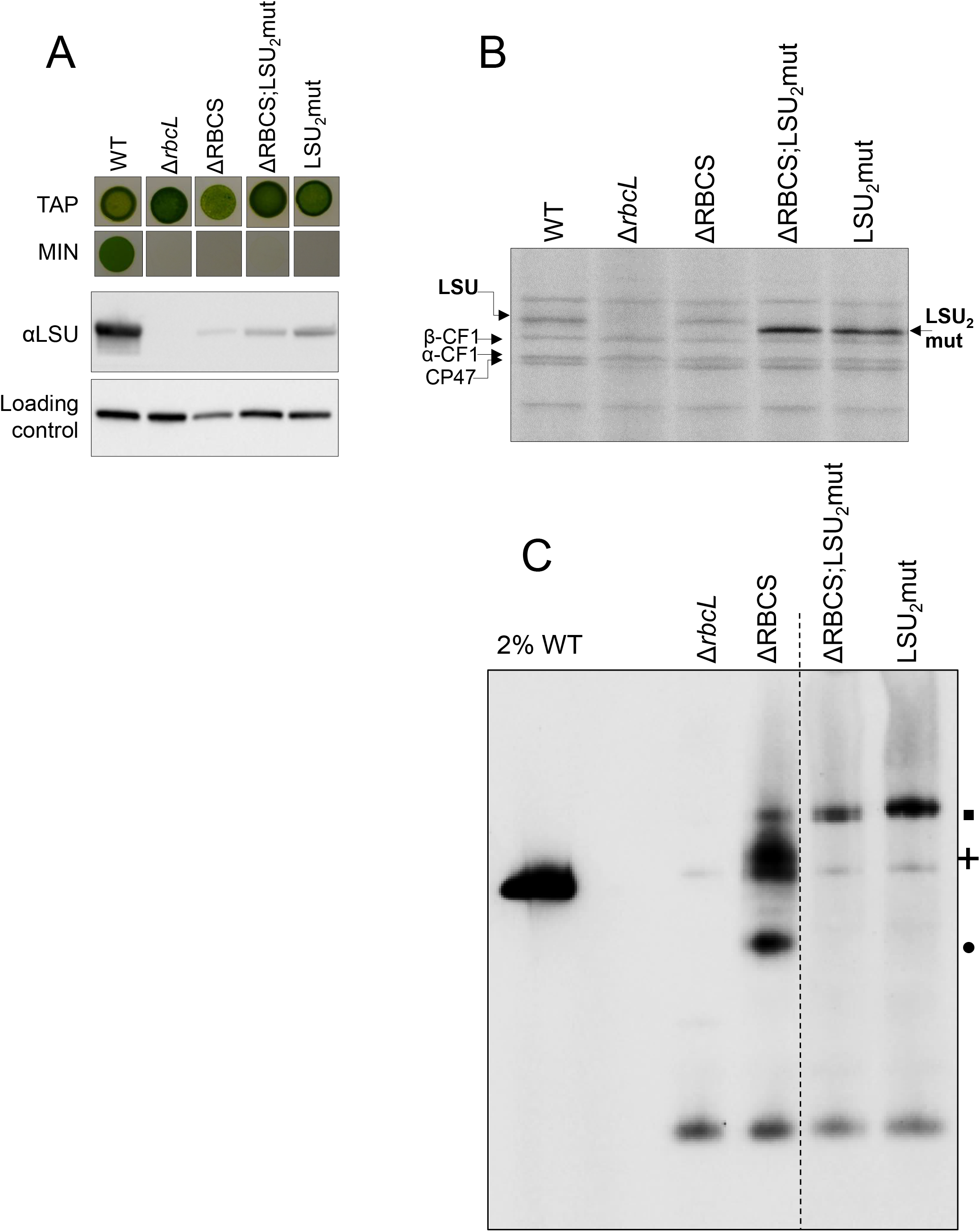
LSU_2_ mutations alter LSU accumulation and CES regulation. (A) Impairment in Rubisco accumulation is revealed by the absence of phototrophic growth in the LSU_2_mut and ΔRBCS;LSU_2_mut strains as probed by spot tests on acetate-free minimal media (MIN). Growth on TAP is shown as a control. The corresponding soluble LSU accumulation detected by immunoblot is shown together with PsaD accumulation as loading control. (B) *RbcL* translation rate in LSU_2_mut and ΔRBCS;LSU_2_mut measured by short ^14^C pulse labeling experiment and compared to ΔRBCS and WT. (C) Immunoblot with the Rubisco antibody after Colorless Native PAGE analysis of soluble protein fractions of WT (note the dilution), ΔrbcL, ΔRBCS, LSU_2_mut and ΔRBCS;LSU_2_mut strains. A dashed line marks the position where two irrelevant lanes were removed. The position of the LSU-complexes observed in ΔRBCS is indicated at the right using the same symbols as in Figure 5 (square, cross and circle).

We then produced another set of mutations in the *rbc*L gene aimed at preserving the ability of LSU to dimerize but not to oligomerize further. According to the 3D structure of Rubisco (PDB 1IR2, (Mizohata et al., 2002)), LSU octameric core shows stabilizing interactions between adjacent LSU dimers involving hydrogen bonds between the guanidino-group of R215 and carbonyls of the main chain of D286 and N287 residues, as well as a salt bridge between the D216 and K161 residues from adjacent LSU dimers. We further noticed that the distance between two LSU dimers was the shortest at the A143 residues which were facing each other closely at the interface of two LSU dimers. To prevent productive interactions between LSU dimers, we introduced a steric clash by replacing the A143 alanine with a bulky tryptophan (A143W, Sup. Fig. 5B) and disrupted the hydrogen bond and salt bridge formation by replacing the arginine and aspartic acid charged residues by neutral ones (R215A-D216A, Sup. Fig. 5B). The resulting triple A143W-R215A-D216A (ARD) substitution was introduced in LSU sequence, yielding LSU_8_mut (rbcL_A143W-R215A-D216A_) transformants, which were produced both in an *RBCS +* (LSU_8_mut) or *RBCS* deficient context (ΔRBCS;LSU_8_mut). As expected and shown for representative strains from the different genotypes in Fig. 9A, the LSU_8_ mutations resulted in a complete loss of phototrophy. Soluble LSU accumulated to levels comparable to that in a ΔRBCS strain, irrespective of the presence or absence of SSU (Fig. 9A). We analyzed LSU translation rate in LSU_8_mut and ΔRBCS;LSU_8_mut by ^14^C pulse-radiolabeling (Fig. 9B). In both cases, LSU was synthesized at a higher rate than WT, irrespective of the presence of SSU (compare LSU_8_mut lane and ΔRBCS; LSU_8_mut lane). Similarly to what had been observed for LSU_2_mut strains, the three substitutions abolished the CES behavior of LSU, allowing its translation to develop in an unregulated configuration.

**Fig. 9:**
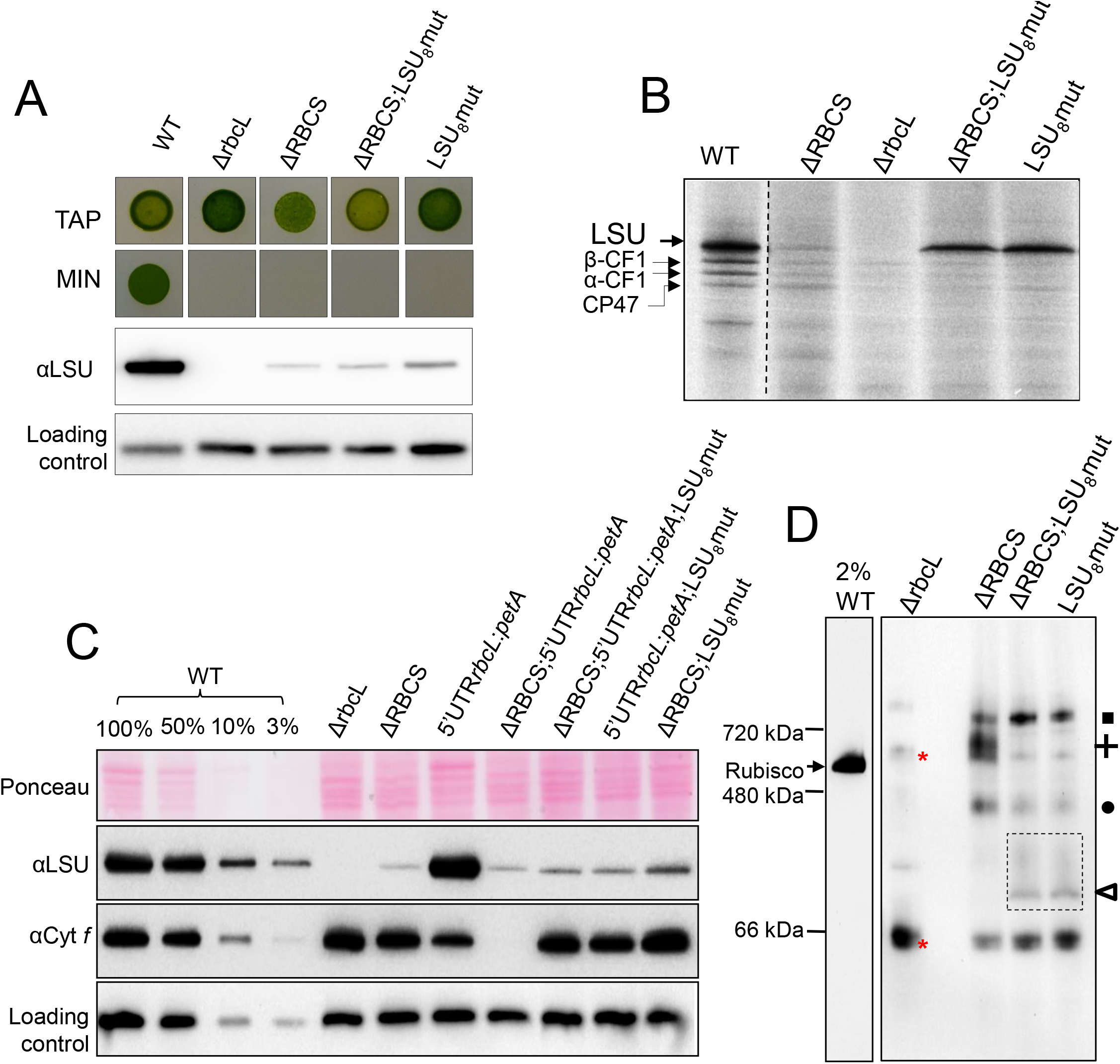
Disruption of LSU oligomerization alters LSU CES regulation. (A) Impairment in Rubisco accumulation is revealed by the absence of phototrophic growth in the LSU_8_mut and ΔRBCS; LSU_8_mut strains as probed by spot tests on acetate-free minimal media (MIN). Growth on TAP is provided as a control. The corresponding soluble LSU accumulation detected by immunoblot is shown. (B) *RbcL* translation rate in LSU_8_mut and ΔRBCS;LSU_8_mut measured by short ^14^C pulse labeling experiment and compared to ΔRBCS and WT. The dashed line marks the position of two irrelevant lanes, which were removed. (C) Immunoblot showing LSU and cytochrome *f* accumulation levels in the wild-type (WT), ΔRBCS, ΔrbcL, LSU_8_mut and ΔRBCS; LSU_8_mut strains and in those latter three genetic contexts combined with the 5’UTRrbcL:*petA* reporter gene background. PsaD accumulation is provided as a loading control. (D) Immunoblot with the Rubisco antibody after Colorless Native PAGE analysis of soluble protein fractions of WT (note the dilution), ΔrbcL, ΔRBCS;LSU_8_mut and ΔRBCS;LSU_8_mut strains. The position of the LSU-complexes observed in ΔRBCS is indicated at the right using the same symbols as in Figure 5 (square, cross, and circle). The empty triangle and dashed box indicate the somewhat diffuse band attributed to LSU dimer.

To further substantiate this conclusion, we combined the same LSU_8_mut substitutions in the presence of the 5’UTRrbcL-*petA* reporter. To this end we introduced the ARD substitutions in a representative ΔRBCS;5’UTRrbcL:*petA* strain, which had undergone *aadA* marker removal (RCalΔK, see Sup. Table 1). These transformants were crossed to the WT strain to segregate the Δ*RBCS* mutation and isolate progenies bearing the LSU_8_ mutations combined to the 5’UTRrbcL-*petA* reporter gene in presence or absence of SSU. We monitored translation of the *petA* reporter by analyzing cyt. *f* accumulation. Fig. 9C shows the results obtained for representative progenies, which should be compared to the original experiment Fig. 3, which demonstrated that the cyt. *f* reporter showed a regular CES regulation when co-expressed with native LSU. In sharp contrast, when co-expressed with LSU bearing the ARD substitutions, the cyt. *f* reporter now accumulated to the same extent, whether SSU was present or not (compare lanes 9 and 10 to lane 8). This shows that synthesis of the cyt. *f* reporter is no longer regulated in the 5’UTRrbcL:*petA*;LSU_8_mut mutants, irrespective of the assembly state of Rubisco. Noteworthy, cytochrome *f* accumulation accumulated to a higher extent when compared to the 5’UTRrbcL-*petA* strain producing native LSU (lane 7). This indicates a higher translation rate of the reporter gene in the CES-insensitive context, and is similar to the increase of LSU synthesis rate observed in the LSU_8_ mutant alone compared to WT. Therefore, when Rubisco assembly can proceed, synthesis does not operate to its maximal rate, neither for LSU, nor for the cytochrome *f* reporter. Altogether, these observations suggest that there is still a significant translation repression in a wild type context for LSU expression.

### Assembly intermediates in the LSU_8_mut oligomerization mutant reveal which LSU form is the CES repressor

The LSU_8_mut mutants, with or without SSU, were analyzed by Native PAGE in comparison with the ΔRBCS strain to characterize the pattern of accumulation of LSU intermediates (Fig. 9D). Detection with the anti-Rubisco antibody yielded a similar pattern for LSU_8_mut and ΔRBCS. The high molecular weight LSU-CPN60 complex observed above 720 kDa and depicted as a square in Fig.9D was present and more abundant than in ΔRBCS strain. The Rubisco-specific band that we attributed to LSU dimer bound to RAF1 (depicted as a circle) was still present, but slightly less abundant. However, a new Rubisco specific band of about 100 kDa, of low abundance and sometimes diffuse appearance, was observed specifically in the two LSU_8_mut mutant strains (Fig. 9D, open triangle). Due to its molecular mass, to the fact that it is not observed in the LSU_2_mut strains and to its absence of immunodetection with a RAF1 antibody (Supp. Fig.6), this band most likely represents a chaperone-free LSU dimer rather than a monomeric LSU form bound to RAF1. Most interestingly, the high molecular weight LSU-RAF1 complex present in ΔRBCS (depicted as a cross in Fig. 5 and 9) was totally absent in the LSU_8_mut mutants. We conclude that the ARD mutations indeed prevented further oligomerization of LSU dimers into the LSU_8_ form, thereby preventing formation of the RAF1-containing HWM-LSU complex, which we tentatively attribute to an LSU_8_-RAF1 species. Thus, this HMW-LSU complex is likely to be the inhibitor of *rbcL* translation in ΔRBCS strain, as its specific disappearance caused by ARD mutations is concurrent with the escape from the CES regulation (Fig. 9 B and C).

## Discussion

### New insights into the pathway for Rubisco biogenesis in vivo

Despite the simple final composition of the hexadecameric (LSU_8_SSU_8_) Rubisco enzyme present in cyanobacteria and chloroplasts of photosynthetic eukaryotes, at least 5 possible partners have been identified in its biogenesis pathway. Besides RBCX, all of these auxiliary proteins were identified in Rubisco-defective mutants (CPN60, (Barkan, 1993); BSD2, (Roth et al., 1996); RAF1, (Feiz et al., 2012), RAF2, (Feiz et al., 2014; Fristedt et al., 2018), indicating their *in vivo* requirement for Rubisco biogenesis in the organism used for the genetic screen. In a few instances, biochemical assays pointed their interaction with LSU and/or SSU ((Barraclough and Ellis, 1980; Feiz et al., 2012; Feiz et al., 2014; Kolesinski et al., 2014). Here we identified *bona fide* assembly intermediates *in vivo* using SSU-defective nuclear mutants and site-directed chloroplast mutants of LSU in Chlamydomonas. Our analysis of a ΔRBCS strain allowed us to detect low amounts of LSU assembly intermediates that would otherwise be obscured by the hundred times more abundant fully assembled holoenzyme. It revealed the existence of two LSU containing species, beside the CPN60-LSU complex previously identified in plants either by *in organello* translation (Roy et al., 1982) or in maize chloroplasts (Feiz et al., 2012). As summarized in Fig. 10, our work supports a pathway where newly-synthesized LSU, arising from the translation of an MRL1-protected *rbcL* mRNA, needs to be kept unfolded, maybe with the help of general chaperones such as a trigger factor (Rohr et al., 2019), until its loading on the CPN60 chaperonin. Its release would be followed by a rapid dimerization, possibly assisted by the RBCX chaperone. RAF1 would subsequently bind, leading to the stabilization of an LSU_2_-RAF1 intermediate. Subsequently, oligomerization would proceed up to an LSU_8_ core, still RAF1-associated, before SSU binding in the ultimate step of Rubisco assembly, thereby displacing the bound chaperone. That SSU requires an LSU octamer for binding, is readily deduced from our observations of (i) a similar accumulation of LSU intermediates, whether SSU is present or not, in the LSU_8_ oligomerization mutant and (ii) a similar pattern of LSU intermediates in the LSU_2_ mutant, irrespective of SSU availability.

**Fig. 10:**
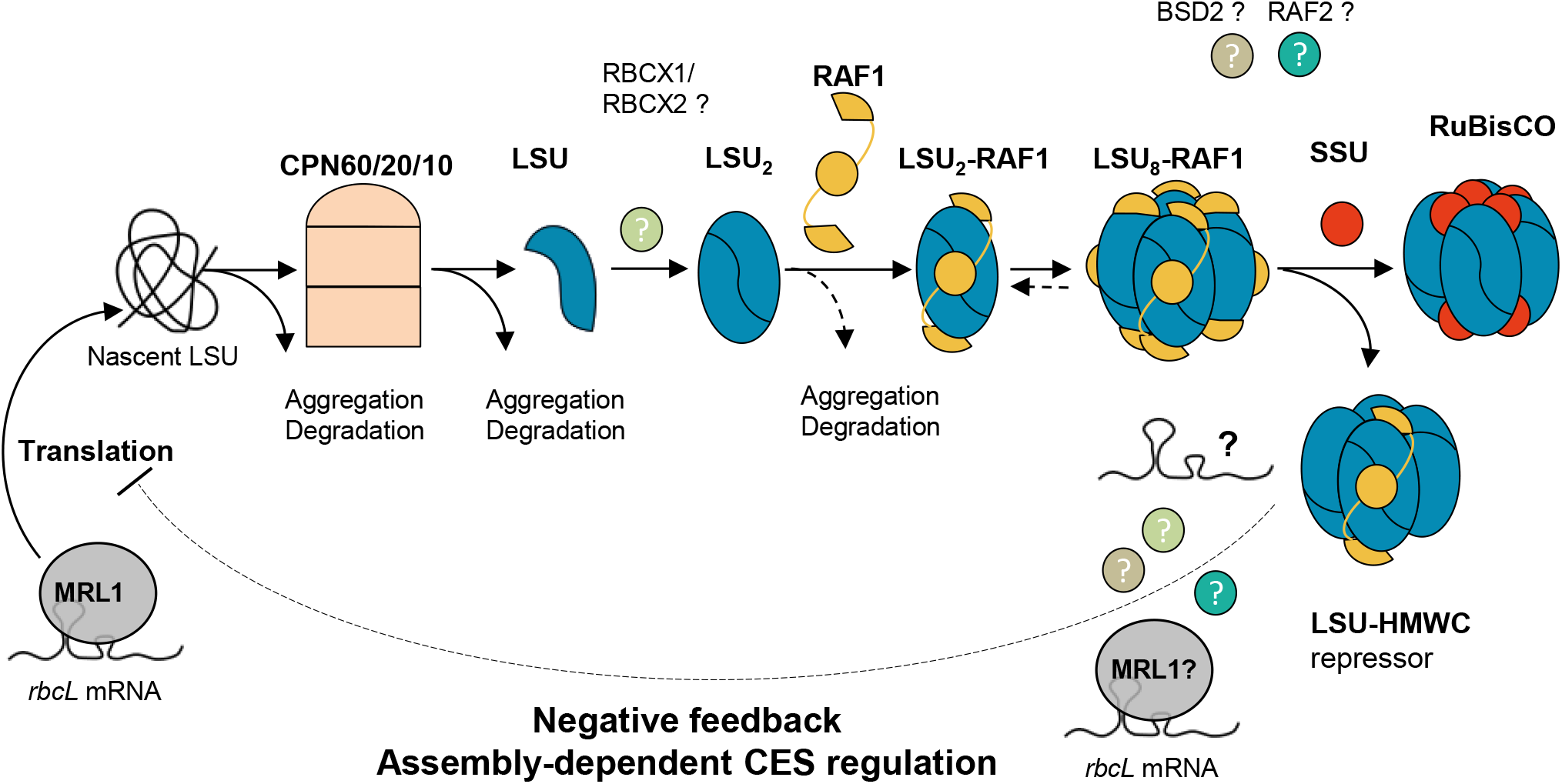
Model of Rubisco biogenesis pathway and CES regulation. The *rbcL* mRNA, stabilized by the binding of the MRL1 PPR-protein to its 5’UTR region, can be translated. Nascent LSU is recruited by the chloroplast folding machinery. LSU propeptide is subsequently folded in the CPN60/20/10 chaperonin complex. The released LSU dimerizes, maybe with help of RBCX, and recruits RAF1 which is required for LSU_2_ stabilization. LSU_2_-RAF1 unit oligomerizes further to form Rubisco catalytic core. RAF1 is finally substituted by the SSU to form the complete holoenzyme. In the SSU-limiting context, the LSU-RAF1 HMWC is converting to a repressor of *rbcL* translation (CES process) preventing LSU wasteful production, by binding either directly *rbcL* mRNA or other factors, thereby displacing some RAF1 oligomers. Many aspects of this model remain unclear such as the identity of the proteins/RNA in the LSU regulator complex, or the exact role of the other Rubisco assembly chaperones such as RBCX1/2 and RAF2, or the presence of a functional homolog of the plant BSD2 factor in algae, which remains debated.

This biogenesis pathway, which results from the present *in vivo* work on the Chlamydomonas enzyme, is further supported by several *in vitro* and *in vivo* studies of the cyanobacterial and plant enzymes (Saschenbrecker et al., 2007; Liu et al., 2010; Bracher et al., 2011; Hauser et al., 2015; Aigner et al., 2017). That RAF1 interacts with LSU in Chlamydomonas (Fig. 6) is consistent with its interaction with LSU in cross-linked maize extracts (Feiz et al., 2012) and in cyanobacteria (Kolesinski et al., 2014; Kolesinski et al., 2017). In agreement with in *vitro* studies of cyanobacterial LSU (Kolesinski et al., 2014; Hauser et al., 2015), we provided several lines of evidence that the RAF1_2_-LSU_2_ dimer, close to 200 kDa (2 × 52 kDa LSU + 2 × 51 kDa RAF1 = 206 kDa), represents indeed a genuine intermediate in the pathway for Rubisco assembly: (i) it was easily detected in a mutant lacking SSU but it was also detected in WT extracts (Fig. 7); (ii) it was absent from an LSU dimerization mutant (Fig. 8C) but it accumulated in absence of further LSU oligomerization, as in the LSU_8_ mutant (Fig. 9D). We note however that LSU dimers without RAF1 still can be formed (Fig. 9D and Sup. Fig. 6). This suggests that RAF1 is not required for the formation of LSU dimers *per se* but rather for their stabilization.

Interestingly, *in organello* pulse-labeling experiments on pea chloroplasts - which have limited availability in unassembled SSU, or chaperones – also identified a 7S LSU associated complex attributed to an LSU dimer (Hubbs and Roy, 1992) and an LSU_8_-like species called Z (Hubbs and Roy, 1993). The size of these two complexes is consistent with that of the LSU oligomers that we identified in the present study (Hubbs and Roy, 1993). Notably, we and others (Hubbs and Roy, 1992; Hubbs and Roy, 1993; Feiz et al., 2012) never detected LSU monomers. These were also absent in the LSU_2_ dimerization mutant despite the enhanced synthesis of LSU in pulse-labeling experiments (Fig. 8). Therefore, LSU monomers are either not proteolytic-resistant or soluble enough to accumulate to detectable levels.

The LSU_8_-RAF1 complex, migrating above the holoenzyme, with an apparent molecular mass of 760 kDa, is present in ΔRBCS, but no longer in the LSU_8_ oligomerization mutant (Fig. 9D). The size of this complex is close to the observed size of the LSU_8_-RAF1_8_ complex identified in reconstitution experiments performed *in vitro* in absence of SSU, using denatured cyanobacterial LSU from *S. elongatus* sp. PCC7942 and RAF1 (Hauser et al., 2015). This LSU_8_-RAF1_8_ complex is believed to constitute the end-point assembly intermediate prior to SSU binding in cyanobacteria. Here, we observed by immunoblotting a higher LSU/RAF1 labeling ratio in the 760 kDa than in the 200 kDa oligomers (Fig. 5). It suggests that there may be as yet unknown interactants in the LSU_8_-RAF1 complex, unless RAF1 is more easily lost from the LSU_8_ core than from LSU_2_ during their purification. The actual composition of the LSU_8_ oligomers before binding of SSU remains confusing. LSU_8_ have been crystallized with RBCX (Bracher et al., 2011), RAF1 (Huang et al., 2020; Xia et al., 2020) and BSD2 (Aigner et al., 2017). However, the last assembly intermediate is considered as a RAF1_8_-LSU_8_ complex in cyanobacteria (Hauser et al., 2015; Kolesinski et al., 2017; Xia et al., 2020), but as a BSD2_8_-LSU_8_ complex in plants (Aigner et al., 2017; Conlan et al., 2019). In view of our results and of the absence of a dedicated BSD2 factor in green algae, we posit that the RAF1-LSU_8_ complex constitutes the penultimate assembly intermediate in green algae. Interestingly, the RAF1-LSU_8_ complex in Chlamydomonas, when probed with an antibody against RAF1, exhibits some change in its migration pattern depending on SSU availability (Fig.6). This raises the possibility of some flexibility in the composition of RAF1-LSU_8_ oligomers depending on the availability for SSU.

### Fate of unassembled LSU

In absence of SSU, which prevents Rubisco assembly, LSU synthesis is inhibited but not fully impaired, as newly-synthesized LSU is readily detected in pulse-labeling experiments (Fig. 1). Yet the LSU accumulation level drops to less than 1%, as estimated by comparing the LSU signal in ΔRBCS to a WT dilution series. We observed that unassembled LSU is stable over more than four hours, as shown by immunochase experiments (Fig. 1D and Sup. Fig. 1B). Our study showed that the discrepancy between a reduced LSU synthesis and a much larger drop in its accumulation in absence of SSU results from both an inhibition of synthesis dependent of the 5’UTR and from the degradation of excess unassembled LSU, independently of the 5’UTR. This is further exemplified by the observation that even with a high rates of synthesis for native LSU, as observed in the ΔRBCS;5’UTRpsaA:*rbcL* strain, the maximal amount of unassembled LSU, accumulating as soluble protein, reaches less than 10 % of the WT level. Altogether, it suggests a bottleneck in LSU assembly, likely due to the limiting amount in one of the assembly chaperones: part of neo-synthesized LSU undergoes rapid proteolysis, when not stabilized by assembly factors. Our data suggest that RAF1 is in limiting amounts. Indeed, we noted that a fraction of LSU dimers is not found associated to RAF1 in the LSU_8_ oligomerization mutant, where LSU is produced in excess, (Fig. 9D and Supp. Fig. 6). That the LSU mutations would rather alter the affinity constant for RAF1 binding, thereby displacing the equilibrium between the RAF-bound and unbound forms of LSU dimers is unlikely since the mutated residues do not belong to the RAF1-LSU interface regions identified in the recently obtained LSU_8_-RAF1_8_ complex crystals (Xia et al., 2020). Rather, this suggests that RAF1 is in limiting amounts, thereby leading to accumulation of LSU dimers without the chaperone. In support to this conclusion, we observed a dramatic increase in the accumulation of unassembled LSU concomitant to RAF1 overexpression using RAF1-epitope tagged strains (compare input fractions of tagged strains versus WT in Fig. 6). Similarly, RAF1 overexpression in maize lines led to a 30% increase in whole Rubisco content (Salesse-Smith et al., 2018).

We further note that the LSU_8_ oligomerization mutant and LSU_2_ dimerization mutant display a similar drop in the accumulation of LSU as in ΔRBCS despite their much higher rates of LSU synthesis. Thus the proteolytic susceptibility of LSU is increased in these mutants (Sup. Fig. 1A and B). One should consider also a possible formation of LSU-insoluble aggregates that would escape recovery in our gels used for electrophoretic purification. Such aggregates have been described by (Cohen et al., 2005) under oxidative stress, or by (Zhan et al., 2015) in the ΔRBCS strain. Although we did not detect such aggregates in ΔRBCS, we did observe some triton-insoluble LSU in the two oligomerization mutants, as well as in the 5’UTRpsaA:*rbcL* expressing strain (data not shown). Whether these result from LSU unproductive interactions prior to LSU loading on the CPN60/CPN23/CPN10 complex, or after its release from the chaperonin is still unknown. We observed in those strains, along with a higher translation rate of the *rbcL* transcript, an increase in the abundance of the CPN60-LSU complex when compared to ΔRBCS (Fig. 9D), suggesting that the chaperonin is not in limiting concentration. Altogether, these observations indicate that there may be several quality checkpoints in Rubisco folding before and after chaperonin interaction, thus directing excess, chaperone-free unassembled LSU subunits either to degradation and/or to aggregation, as depicted in figure 10.

### The CES process for LSU

An efficient biogenesis of the oligomeric proteins which build up the photosynthesis apparatus requires some coordination in the expression of their subunits. It is even more so when their subunits are encoded in distinct intracellular compartments, which precludes a transcriptional co-regulation as is the case in photosynthetic bacteria. Indeed, we previously demonstrated that a number of chloroplast genes encoding core subunits from photosynthetic complexes in Chlamydomonas, undergo regulatory loops depending on their assembly state with the other subunits of the same protein complex. This feedback, called the CES process (Control by Epistasy of synthesis) (Wollman et al., 1999; Choquet and Wollman, 2007), occurs at the level of translation initiation. Its importance is reflected by its prevalence, as CES subunits have been identified in all membrane-embedded photosynthetic proteins, Photosystems I and II, cyt *b*_*6*_*f* and ATP synthase, allowing a fine-tuning of their expression by the presence of their assembly partners. Here we showed that Rubisco also displays a CES behavior for its biogenesis in Chlamydomonas. Rubisco is a most interesting case, as a CES behavior for LSU also has been observed in higher plant chloroplasts (Wostrikoff and Stern, 2007), providing the first example of the conservation of this regulatory process in multicellular eukaryotes. Chlamydomonas offers unique opportunities to shed more light on the CES mechanism for Rubisco because it is amenable to genetic approaches. This allowed us to perform experiments with Chlamydomonas, which are presently not feasible in plants, to provide a complete demonstration of the control of *rbcL* expression by the state of LSU assembly.

We first showed that the CES process for LSU synthesis is exerted at the level of initiation of translation. We came to this conclusion after swapping *rbcL* 5’UTR. We demonstrated that this gene region contains all cis-elements required for the assembly-dependent regulation of *rbcL* translation. This excludes both an effect on translation elongation and an early co-translational degradation of LSU, both of which would target its coding sequence and not the untranslated region of the gene. This is a feature which distinguishes the CES process from the other known translational regulation of LSU, which were attributed to a change in its rate of translational elongation such as LSU inhibition of translation elongation in the dark (Mühlbauer and Eichacker, 1998) and LSU translational repression under oxidative stress (Shapira et al., 1997). In the latter case, a structural conformational modification in oxidized LSU was suggested to expose an otherwise buried N-terminal domain. This domain adopts a ferredoxin fold structure, similar to an RNA Binding domain (RBD), and was found to have an unspecific RNA binding capacity (Yosef et al., 2004; Cohen et al., 2005).

As the CES process results from an autoregulation of translation by LSU, it should be mediated by either one of the three stable assembly intermediate found to accumulate in absence of SSU. The first step of LSU folding, CPN60 bound LSU, does not serve as a regulator of translation since the two oligomerization mutants that we tested accumulated higher level of this intermediate but showed no translational repression despite the absence of Rubisco assembly. In contrast, the LSU_8_-RAF1 complex displays the properties expected for a negative-feedback regulator: its absence correlates with the escape from the CES process in both the dimerization and LSU_8_mut mutants (*rbcL*_E109A-R253A_ and *rbcL*_A143W-R215A-D216A_). Interestingly, our data also ruled out the possibility that the repressor would be constituted by free monomeric LSU or by the LSU dimer which is not compromised in the LSU_8_mut mutant. Thus, the LSU octamer accumulated in absence of the SSU partner is the key LSU form which provides the assembly-dependent translational control.

Whether RAF1 plays an active role in the regulation of *rbcL* translation, or only mediates the formation of the HMW-LSU repressor form remains to be determined. Notably, in maize, where LSU is also under the CES regulation (Wostrikoff et al., 2012), the RAF1 knock-out mutant displays a high LSU synthesis rate (Feiz et al., 2012). This observation is compatible with RAF1 being required for the formation of the translational repressor, but does not allow to reveal RAF1’s role in the repression. Facts detracting RAF1 direct involvement in this process come from its unexpected stoichiometry to LSU. While we could not retrieve from our data the exact stoichiometry of RAF1 to LSU in the repressor form, it seems unlikely that the repressor complex comprises 8 LSU subunits bound to 4 RAF1 dimers, as the RAF1 to LSU ratio is decreased in the LSU_8_-RAF1 complex compared to the LSU_2_-RAF1 dimer. This is at variance with cyanobacterial reconstitution experiments that identify the LSU_8_-RAF1_8_ complex to be stable (Hauser et al., 2015). As noted above, there is a slight but significant difference in migration of the HMW-RAF1 complex in the WT compared to the ΔSSU strain (Fig. 7). Additional experiments on this issue are challenged by the presence of large amounts of Rubisco complexes overlapping with LSU_8_-RAF1 in the wild type. It is tempting to consider that it corresponds to the LSU_8_-RAF1_8_ assembly intermediate, which has been observed and crystallized in cyanobacteria (Hauser et al., 2015; Xia et al., 2020), which would convert to a repressor form complex in absence of SSU, leading to this slight change in migration in our gel system. The observed upshift may result from the capture of other proteins or RNA thereby leading to translational repression (Fig. 10).

The molecular mechanism by which LSU_8_-RAF1 controls translation of the *rbcL* transcript, could result from the RNA binding activity of Rubisco RNA Recognition Motif that would allow the *rbcL* mRNA to get sequestered in this complex, thereby rendering it unavailable for translation. In such a case, repression would result from a direct interaction between LSU and its transcript. Whether the RRM domain is indeed accessible in the repressor conformation, cannot be reasonably addressed without better knowledge of the components of this repressor form which is a challenge owing to its very low accumulation level. Alternatively, CES translation inhibition could be mediated by an additional trans-acting factor, as has been documented for the CES process governing cyt. *f* translation. In this model, the MCA1 protein that is responsible for *petA* mRNA stabilization and translation is targeted to degradation via its interaction with a repressor motif exposed when cyt. *f* remains unassembled (Boulouis et al., 2011). The MRL1 factor is an obvious candidate for this function: it interacts with *rbcL* mRNA and promotes its stabilization. Furthermore MRL1 is found in a large complex, whose size is dependent upon LSU presence (Johnson et al., 2010). Yet, the proposed mechanism would be different as we found MRL1 to be stable at variance with MCA1 (data not shown). MRL1 could be part of the LSU repressor complex, thereby leading to *rbcL* transcript sequestration away from the ribosomes. However, in this model the absence of MRL1 should alter the migration pattern of the LSU_8_-RAF1 complex on native gels, which is not observed (Suppl. Fig. 7). Alternatively, an as for now undetected interactant, such as RBCX, could indirectly mediate this interaction. Interestingly, a similar model has recently been proposed to be involved in D1 translation in higher plant chloroplasts. The presence of a D1-HCF244-OHP1/2 assembly intermediate was linked to the inhibition of D1 synthesis in the dark, relieved in the photorepair process. This assembly intermediate has been suggested to act as a repressor complex which may physically interact with D1 HCF73 translational activator to mediate D1 repression in the dark (Chotewutmontri and Barkan, 2020).

In conclusion, our suggestion that the repressor form identified in absence of assembly constitutes a regulatory pool, rather than representing a true end-state assembly intermediate in Chlamydomonas, further raises the question whether it would participate in regulating LSU synthesis rates in regular WT conditions. The large amount of assembled Rubisco precludes us from testing whether the same LSU_8_-RAF1 complex is present in WT. However the observation that LSU synthesis rate may be higher than what is observed in WT conditions, as observed in the dimerization or oligomerization mutants, hints to the fact that LSU synthesis is probably limited by a CES mechanism even in WT, as it has been reported for cytochrome *f* (Choquet et al., 1998; Choquet and Wollman, 2007). Further experiments are required to determine the exact composition of the LSU_8_-RAF1 complex in order to precise the exact mechanism of the assembly-mediated *rbcL* translation inhibition and confirm its occurrence or not in WT conditions.

## Materials and Methods

### Cultures and strains

If not stated otherwise, wild type (WT222 mt+ and WTS24 mt-) and mutant strains of *C. reinhardtii* were grown on solid Tris-acetate-phosphate medium (TAP) (Harris, 2009) supplemented with agar and in liquid cultures under continuous, dim light (7 μmol photons.m^−2^.s^−1^) on an orbital shaker (120 rpm) at 25°C. Cells from exponentially growing cultures (2 × 10^6^ cells.mL^−1^) were used for all experiments.

### Chlamydomonas genetics methods

Chloroplast transformation was done as described in (Kuras and Wollman, 1994) using an in house built helium-driven particle gun. Chlamydomonas mating and progeny isolation were done as in (Harris, 2009).

### Nucleic acids manipulations

If not stated otherwise DNA manipulations were done following standard protocols as in (Sambrook et al., 1989). RNA extractions and blotting was performed as in (Drapier et al., 2002).

### Plasmids and strains preparation

Plasmids carrying mutations aimed to introduce a truncation (pLStr) or a triple ARD substitution in LSU sequence (pLS_ARD_), to prevent LSU dimerization (pL_2_mut), or carrying the *psaA*-driven *rbcL* gene (paAR) are described in supplemental Materials and Methods. All plasmids contain the *5’psaA-aadA-atpB* 3’ selection marker conferring resistance to spectinomycin, flanked by direct repeats (Fischer et al., 1996) allowing the cassette removal in absence of selection pressure, at neutral positions either at *rbcL* 5’ (*Bse*RI site) or 3’end (*Afl*II site). Plasmid pRFFFiK aimed to express the *petA* gene from *rbcL* 5’ regulatory regions was described in (Johnson et al., 2010). Biolistic transformation was used to transform appropriate recipient strains, which were as specified in the supplemental table 1 the WT T222+ strain, the *rbcL* deletion strain ΔR T1.3+, the *RBCS* mutant Cal.13.5A+ strain (back-crossed progeny of the CAL005.01.13 strain described in (Dent et al., 2005), and RCalΔK strain (Cal13.5A+ transformed with pRFFFiK (containing the *petA* sequence where the endogenous *petA* 5’UTR was swapped by the *rbcL* promoter and 5’UTR, described in (Johnson et al., 2010)) and subjected to cassette removal (Fischer et al., 1996)). Transformants were brought to homoplasmy by 6 rounds of subcloning on selective media (TAP supplemented with 500ug/mL Spectinomycin), after which homoplasmy was confirmed by PCR analysis.

Cloning of the pJHL-Raf1S plasmid, allowing expression of the RAF1 gene fused at its C-terminus to a StrepII tag driven by the PsaD promoter and carrying the *aphVIII* resistance gene, is described in supplemental Materials and Methods. It was used to transform the ΔRBCS strain. Nuclear transformation was performed according to (Onishi and Pringle, 2016), and transformants selected on TAP supplemented with 10 ug/L paromomycin.

### Pulse experiment

Chlamydomonas ^14^C-acetate pulse experiment was done as described in (Drapier et al., 2007). 5 × 10^6^ cells were washed once in MIN-Tris medium then resuspended in 5 ml fresh MIN– Tris medium and incubated for 1h at RT with vigorous shaking to remove the acetate from the medium at a light intensity of 30 μmol photons.m^−2^.s^−1^. Subsequently, 5 μl of 1mg.ml^−1^ cycloheximide and 50 μCie of ^14^C-acetate were added simultaneously to the cells. After 7 min of vigorous shaking cells were mixed with 35 ml of cold TAP medium supplemented with 40mM acetate and immediately spun down. Cell pellets were afterwards washed in 5 mM Hepes buffer supplemented with EDTA-free protease inhibitors mix (Roche), resuspended in 0.1M DTT, 0.2M Na_2_CO_3_ and flash-frozen. Prior to denaturation, samples were suspended 1:1 in 5% SDS, 20% sucrose solution, boiled for 1 minute, then spun down at 12 000g for 15 minutes. The supernatant was subsequently loaded on urea 12-18% polyacrylamide gradient gels using in house-built gel system. Samples from pulse labeling experiments relying on ^14^C incorporation in the alga are loaded based on radioactivity incorporation rather than equal protein amount and rates of synthesis are estimated by comparison to labeling intensity of unrelated chloroplast translates. After migration, gels were stained with Coomassie Blue, dried and exposed to an autoradiography screen for at least one month. Phosphorescence signal was measured using a Typhoon FLA 9500 phosphorimager (GE Healthcare).

### Immunochase experiments

Exponentially growing Chlamydomonas cells from a 200 mL culture were treated with chloramphenicol at a final concentration of 500 μg/ml. 30 mL aliquots were removed along the chase at the initial time-point of inhibitor addition and after 1h, 2h and 4h, and were submitted to protein extraction followed by separation on 12% SDS-PAGE. Samples from different time-points were loaded on an equal chlorophyll basis, as in (Kuras and Wollman, 1994).

### Protein analysis

*Protein electrophoresis* in denaturing conditions was performed according to the modified Laemmli protocol (Laemmli, 1970). For protein loading of “whole cell” samples, chlorophyll fluorescence was used for quantification as in (Kuras and Wollman, 1994). Samples were suspended 1:1 in 5% SDS, 20% sucrose solution and boiled for 1 minute, then spun down at 12.000g for 15 minutes to remove insoluble material and subsequently analyzed using 12% (in-house gels) or 8-16% SDS-polyacrylamide gels (Bio-Rad).

*Colorless Native electrophoresis* (CN-PAGE) was done according to a modified Schägger protocol (Wittig and Schagger, 2005). Cell pellets from 200 mL of Chlamydomonas culture were resuspended in an extraction buffer (20 mM HEPES pH= 7.5, 20 mM KCl, 10% glycerol, 2× EDTA-free protease inhibitor mix (ROCHE)), and broken using a French press apparatus at 6 .000 psi. The soluble fraction was collected after centrifugation at 267.000 rcf at 4°C for 25 min and concentrated using Amicon Ultra centrifugation units with a 30 kDa cutoff (Millipore). Protein concentrations were measured colorimetrically by Bradford assay (Bradford, 1976) using QuickStart Bradford Dye reagent (Bio-Rad). 70 μg of protein were loaded on MiniProtean 4-16% gradient gels (Invitrogen), as well as a native protein marker (NativeMark unstained Protein standard, Life Technologies). Migration was undertaken at 4°C at constant voltage of 60 V for 1h than 120 V. Native gels used for immunoblotting were first incubated 1 hour in 2% SDS, 0,67% β-mercaptoethanol prior to their transfer on nitrocellulose membranes (2h30 at 1mA per cm^2^).

*Coimmunoprecipitation* was performed using 400 ug of soluble extracts, prepared as specified in the previous section (NEB buffer: Hepes-KOH pH7.5 20mM, KCl 100mM, glycerol 10%), which were incubated for 30 min at 4°C with 50 ul of ferrimagnetic StrepTactin beads (MagStrep Type3 XT beads, IBA LifeSciences) preincubated in 0.1M Tris pH8.0, 150 mM NaCl, with regular mixing. A fraction of soluble extract (1/8 of the initial amount) was set aside to evaluate the protein composition of the input fraction. A magnetic stand was used to clear the beads from the supernatant. The unbound extract was then removed, and an equivalent fraction (volume wise) set aside. 3X Laemmli loading buffer was added to both the input and unbound fraction. Beads were washed three times in wash buffer (0.1M Tris pH8.0, 150 mM NaCl supplemented with protease inhibitors), and then resuspended in 1X Laemmli loading Buffer. Equivalent fractions of the input, unbound and eluted fractions were denatured for 1 min at 100°C, cleared by centrifugation at 13 000 rpm at 4°C, and separated on a 10% acrylamide-urea 8M-SDS gel, followed by blotting and immunodecoration as specified in the next section.

For *immunoblot analysis*, proteins were transferred onto nitrocellulose membranes (0.1 μ pore size, Amersham Protran). The membranes were blocked with 5% (w/v) skim milk in a phosphate-buffered saline (PBS) solution plus 0.1% (w/v) Tween 20 (Sigma-Aldrich). The target proteins were immunodecorated with primary antibodies and then incubated with horseradish peroxidase-conjugated anti-rabbit IgG antibodies (Catalogue number: W4011, Promega) at a dilution ratio of 1:20.000. Primary antibodies used were directed against Rubisco whole holoenzyme (kindly provided by Dr. Spencer Whitney, used at a dilution of 1:40000 to 1:80.000), Cpn60α/β1 (kind gift of Michael Schroda, used at a dilution of 1:2.000). Antibodies directed against the PSI subunit PsaD and α -tubulin were purchased from Agrisera (catalogue number: AS09 461 and AS10 680) and used respectively at a dilution of 1:10.000 and 1:10.000). Antibodies against cytochrome *f* (PetA; used at a dilution ratio of 1:100.000) were prepared using purified cytochrome *f* injected to rabbits. For RAF1 antibody production, recombinant *C. reinhardtii* RAF1 (Cre06.g308450) was expressed in *E.coli* using a codon-adapted synthetic cDNA (Genscript, Piscataway, NJ, USA). The protein was purified using GST-tag affinity and used directly as an antigen in rabbits (Genscript, Piscataway, NJ, USA). The resulting antiserum was used at a dilution of 1:30.000. Immuno-reactive proteins were detected with Clarity One detection reagent (Bio-Rad) and visualized using the ChemiDoc XRS+ System (Bio-Rad).

## Supplemental Data files

- Supplemental Materials and Methods: plasmid construction and primer list.
- Supplemental table 1: Summary of transformation experiments
- Supplemental figure 1: LSU stability and estimated half-life in different Rubisco mutants.
- Supplemental figure 2: Truncated LSU does not accumulate
- Supplemental figure 3: Anti-RAF1 antibody
- Supplemental figure 4: RAF1 accumulates independently of Rubisco.
- Supplemental figure 5: Close-up of the mutated residues in LSU structure
- Supplemental figure 6: LSU dimers without RAF1 are detected in the octamerization LSU_8_mut mutant
- Supplemental figure 7: The LSU-HMW complex is not affected by MRL1 absence in the *mrl1;ΔRBCS;5’UTR psaA*:rbcL strain

## Acknowledgements

We thank Sonia Fieulaine (I2BC, France) for her kind help in the design of the structure guided mutagenesis approach. We thank Michael Schroda (University of Kaiserslautern, Germany) and Spencer M. Whitney (ANU Canberra, Australia) for kindly providing the anti-CPN60 and anti-Rubisco antibodies respectively. We are grateful to Wojciech Majeran for sharing unpublished results and to Olivier Vallon for providing the Rubisco small subunit mutant strains. We acknowledge basic support from the Centre National de la Recherche Scientifique (CNRS) and Sorbonne Université (SU) to UMR7141, and competitive funding from Labex Dynamo (ANR-11-LABX-0011-01). W.W. benefited from a doctoral support from ED515, Complexité du Vivant and Labex Dynamo (ANR-11-LABX-0011-01). We thank David Stern (BTI, Ithaca, NY) and PICS 5462 for support to K.W. during the initial stage of this project. Finally we thank anonymous reviewers for their insightful comments that helped improve this manuscript.

## Authors contribution

W.W., F.A.W and K.W. designed the research; W.W., E.T. and K.W. performed research; W.W., F.A.W and K.W analyzed data and wrote the paper.

## Bibliography

Aigner, H., Wilson, R.H., Bracher, A., Calisse, L., Bhat, J.Y., Hartl, F.U., and Hayer-Hartl, M. (2017). Plant RuBisCo assembly in *E. coli* with five chloroplast chaperones including BSD2. Science 358, 1272–1278.

Andersson, I., and Backlund, A. (2008). Structure and function of Rubisco. Plant Physiol. Biochem. 46, 275–291.

Bar-On, Y.M., and Milo, R. (2019). The global mass and average rate of rubisco. Proc. Natl. Acad. Sci. USA 116, 4738–4743.

Barkan, A. (1993). Nuclear Mutants of Maize with Defects in Chloroplast Polysome Assembly Have Altered Chloroplast RNA Metabolism. Plant Cell 5, 389–402.

Barkan, A., and Small, I. (2014). Pentatricopeptide repeat proteins in plants. Annu. Rev. Plant Biol. 65, 415–442.

Barraclough, R., and Ellis, R.J. (1980). Protein synthesis in chloroplasts. IX. Assembly of newly-synthesized large subunits into ribulose bisphosphate carboxylase in isolated intact pea chloroplasts. Biochim. Biophys. Acta 608, 19–31.

Börner, T. (2017). The discovery of plastid-to-nucleus retrograde signaling—a personal perspective. Protoplasma 254, 1845–1855.

Boulouis, A., Raynaud, C., Bujaldon, S., Aznar, A., Wollman, F.A., and Choquet, Y. (2011). The Nucleus-Encoded trans-Acting Factor MCA1 Plays a Critical Role in the Regulation of Cytochrome f Synthesis in Chlamydomonas Chloroplasts. The Plant Cell 23, 333–349.

Bracher, A., Starling-Windhof, A., Hartl, F.U., and Hayer-Hartl, M. (2011). Crystal structure of a chaperone-bound assembly intermediate of form I Rubisco. Nat. Struct. Mol. Biol. 18, 875–880.

Bracher, A., Whitney, S.M., Hartl, F.U., and Hayer-Hartl, M. (2017). Biogenesis and Metabolic Maintenance of Rubisco. Annual Review of Plant Biology 68, 29–60.

Bracher, A., Hauser, T., Liu, C., Hartl, F.U., and Hayer-Hartl, M. (2015). Structural Analysis of the Rubisco-Assembly Chaperone RbcX-II from Chlamydomonas reinhardtii. PloS one 10, 1–17.

Bradford, M.M. (1976). A rapid and sensitive method for the quantitation of microgram quantities of protein utilizing the principle of protein-dye binding. Anal. Biochem. 72, 248–254.

Brutnell, T.P., Sawers, R.J., Mant, A., and Langdale, J.A. (1999). BUNDLE SHEATH DEFECTIVE2, a novel protein required for post-translational regulation of the rbcL gene of maize. The Plant Cell 11, 849–864.

Busch, F.A., Tominaga, J., Muroya, M., Shirakami, N., Takahashi, S., Yamori, W., Kitaoka, T., Milward, S.E., Nishimura, K., Matsunami, E., Toda, Y., Higuchi, C., Muranaka, A., Takami, T., Watanabe, S., Kinoshita, T., Sakamoto, W., Sakamoto, A., and Shimada, H. (2020). Overexpression of BUNDLE SHEATH DEFECTIVE 2 improves the efficiency of photosynthesis and growth in Arabidopsis. Plant J. 102, 129–137.

Chaddock, A.M., Mant, A., Karnauchov, I., Brink, S., Herrmann, R.G., Klosgen, R.B., and Robinson, C. (1995). A new type of signal peptide: central role of a twin-arginine motif in transfer signals for the delta pH-dependent thylakoidal protein translocase. EMBO J. 14, 2715–2722.

Chan, K.X., Phua, S.Y., Crisp, P., McQuinn, R., and Pogson, B.J. (2016). Learning the Languages of the Chloroplast: Retrograde Signaling and Beyond. Annual Review of Plant Biology 67, 25–53.

Checa, S.K., and Viale, A.M. (1997). The 70-kDa heat-shock protein/DnaK chaperone system is required for the productive folding of ribulose-biphosphate carboxylase subunits in Escherichia coli. Eur. J. Biochem. 248, 848–855.

Chen, X., Kindle, K.L., and Stern, D.B. (1995). The initiation codon determines the efficiency but not the site of translation initiation in Chlamydomonas chloroplasts. Plant Cell 7, 1295–1305.

Choquet, Y., and Wollman, F.-A. (2007). The CES process. (Academic Press).

Choquet, Y., Stern, D.B., Wostrikoff, K., Kuras, R., Girard-Bascou, J., and Wollman, F.A. (1998). Translation of cytochrome f is autoregulated through the 5′ untranslated region of petA mRNA in Chlamydomonas chloroplasts. Proc. Natl. Acad. Sci. USA 95, 4380–4385.

Chotewutmontri, P., and Barkan, A. (2020). Light-induced *psbA* translation in plants is triggered by photosystem II damage via an assembly-linked autoregulatory circuit. Proc. Nat. Acad. Sci. USA 117, 21775–21784.

Cohen, I., Knopf, J.A., Irihimovitch, V., and Shapira, M. (2005). A Proposed Mechanism for the Inhibitory Effects of Oxidative Stress on Rubisco Assembly and Its Subunit Expression. Plant Phys. 137, 738–746.

Conlan, B., Birch, R., Kelso, C., Holland, S., De Souza, A.P., Long, S.P., Beck, J.L., and Whitney, S.M. (2019). BSD2 is a Rubisco-specific assembly chaperone, forms intermediary hetero-oligomeric complexes, and is nonlimiting to growth in tobacco. Plant Cell Environ. 42, 1287–1301.

Dent, R.M., Haglund, C.M., Chin, B.L., Kobayashi, M.C., and Niyogi, K.K. (2005). Functional genomics of eukaryotic photosynthesis using insertional mutagenesis of *Chlamydomonas reinhardtii*. Plant Phys. 137, 545–556.

Doron, L., Goloubinoff, P., and Shapira, M. (2018). ZnJ2 Is a Member of a Large Chaperone Family in the Chloroplast of Photosynthetic Organisms that Features a DnaJ-Like Zn-Finger Domain. Front Mol. Biosci. 5, 2.

Doron, L., Segal, N., Gibori, H., and Shapira, M. (2014). The BSD2 ortholog in *Chlamydomonas reinhardtii* is a polysome-associated chaperone that co-migrates on sucrose gradients with the rbcL transcript encoding the Rubisco large subunit. Plant J. 80, 345–355.

Drapier, D., Girard-Bascou, J., Stern, D.B., and Wollman, F.A. (2002). A dominant nuclear mutation in Chlamydomonas identifies a factor controlling chloroplast mRNA stability by acting on the coding region of the atpA transcript. Plant J. 31, 687–697.

Drapier, D., Rimbault, B., Vallon, O., Wollman, F.-A., and Choquet, Y. (2007). Intertwined translational regulations set uneven stoichiometry of chloroplast ATP synthase subunits. EMBO J. 26, 3581–3591.

Ellis, R.J. (1979). The most abundant protein in the world. TiBS 4, 241–244.

Emlyn-Jones, D., Woodger, F.J., Price, G.D., and Whitney, S.M. (2006). RbcX can function as a rubisco chaperonin, but is non-essential in Synechococcus PCC7942. Plant Cell Physiol. 47, 1630–1640.

Feiz, L., Williams-Carrier, R., Wostrikoff, K., Belcher, S., Barkan, A., and Stern, D.B. (2012). Ribulose-1,5-bis-phosphate carboxylase/oxygenase accumulation factor 1 is required for holoenzyme assembly in maize. The Plant Cell 24, 3435–3446.

Feiz, L., Williams-Carrier, R., Belcher, S., Montano, M., Barkan, A., and Stern, D.B. (2014). A protein with an inactive pterin-4a-carbinolamine dehydratase domain is required for Rubisco biogenesis in plants. Plant J. 80, 862–869.

Fischer, N., Stampacchia, O., Redding, K., and Rochaix, J.-D. (1996). Selectable marker recycling in the chloroplast. M.G.G. 251, 373–380.

Fristedt, R., Hu, C., Wheatley, N., Roy, L.M., Wachter, R.M., Savage, L., Harbinson, J., Kramer, D.M., Merchant, S.S., Yeates, T., and Croce, R. (2018). RAF2 is a RuBisCO assembly factor in Arabidopsis thaliana. Plant J. 94, 146–156.

Grimm, R., Grimm, M., Eckerskorn, C., Pohlmeyer, K., Röhl, T., and Soll, J. (1997). Postimport methylation of the small subunit of ribulose-1,5-bisphosphate carboxylase in chloroplasts. FEBS Letters 408, 350–354.

Hammani, K., Bonnard, G., Bouchoucha, A., Gobert, A., Pinker, F., Salinas, T., and Giegé, P. (2014). Helical repeats modular proteins are major players for organelle gene expression. Biochimie 100, 141–150.

Harris, E.H. (2009). Chlamydomonas in the laboratory. In The Chlamydomonas Sourcebook (Second Edition) (London: Academic Press), pp. 241–302.

Hartl, F.U. (1996). Molecular chaperones in cellular protein folding. Nature 381, 571–579.

Hauser, T., Bhat, J.Y., Milicic, G., Wendler, P., Hartl, F.U., Bracher, A., and Hayer-Hartl, M. (2015). Structure and mechanism of the Rubisco-assembly chaperone Raf1. Nat. Struct. Mol. Biol. 22, 720–728.

Houtz, R.L., Magnani, R., Nayak, N.R., and Dirk, L.M. (2008). Co- and post-translational modifications in Rubisco: unanswered questions. J. Exp. Bot. 59, 1635–1645.

Huang, F., Vasieva, O., Sun, Y., Faulkner, M., Dykes, G.F., Zhao, Z., and Liu, L.-N. (2019). Roles of RbcX in Carboxysome Biosynthesis in the Cyanobacterium *Synechococcus elongatus* PCC7942. Plant Phys. 179, 184–194.

Huang, F., Kong, W.-W., Sun, Y., Chen, T., Dykes, G.F., Jiang, Y.-L., and Liu, L.-N. (2020). Rubisco accumulation factor 1 (Raf1) plays essential roles in mediating Rubisco assembly and carboxysome biogenesis. Proc. Natl. Acad. Sci. USA 117, 17418–17428.

Hubbs, A., and Roy, H. (1992). Synthesis and Assembly of Large Subunits into Ribulose Bisphosphate Carboxylase/Oxygenase in Chloroplast Extracts. Plant Phys. 100, 272–281.

Hubbs, A.E., and Roy, H. (1993). Assembly of in vitro synthesized large subunits into ribulose-bisphosphate carboxylase/oxygenase. Formation and discharge of an L8-like species. J. Biol. Chem. 18, 6–10.

Jarvis, P. (2008). Targeting of nucleus-encoded proteins to chloroplasts in plants. New Phytologist 179, 257–285.

Johnson, X., Wostrikoff, K., Finazzi, G., Kuras, R., Schwarz, C., Bujaldon, S., Nickelsen, J., Stern, D.B., Wollman, F.A., and Vallon, O. (2010). MRL1, a conserved Pentatricopeptide repeat protein, is required for stabilization of rbcL mRNA in Chlamydomonas and Arabidopsis. The Plant Cell 22, 234–248.

Khrebtukova, I., and Spreitzer, R.J. (1996). Elimination of the Chlamydomonas gene family that encodes the small subunit of ribulose-1,5-bisphosphate carboxylase/oxygenase. Proc. Natl. Acad. Sci. USA 93, 13689–13693.

Kolesinski, P., Piechota, J., and Szczepaniak, A. (2011). Initial characteristics of RbcX proteins from *Arabidopsis thaliana*. Plant Mol. Biol. 77, 447–459.

Kolesinski, P., Rydzy, M., and Szczepaniak, A. (2017). Is RAF1 protein from Synechocystis sp. PCC 6803 really needed in the cyanobacterial Rubisco assembly process? Photosynth. Res. 132, 135–148.

Kolesinski, P., Belusiak, I., Czarnocki-Cieciura, M., and Szczepaniak, A. (2014). Rubisco Accumulation Factor 1 from *Thermosynechococcus elongatus* participates in the final stages of ribulose-1,5-bisphosphate carboxylase/oxygenase assembly in *Escherichia coli* cells and in vitro. FEBS J. 281, 3920–3932.

Kolesinski, P., Golik, P., Grudnik, P., Piechota, J., Markiewicz, M., Tarnawski, M., Dubin, G., and Szczepaniak, A. (2013). Insights into eukaryotic Rubisco assembly - crystal structures of RbcX chaperones from *Arabidopsis thaliana*. Biochim. Biophys. Acta 1830, 2899–2906.

Kuras, R., and Wollman, F.A. (1994). The assembly of cytochrome b6/f complexes: an approach using genetic transformation of the green alga *Chlamydomonas reinhardtii*. EMBO J. 13, 1019–1027.

Laemmli, U.K. (1970). Cleavage of structural proteins during the assembly of the head of bacteriophage T4. Nature 227, 680–685.

Langer, T., Lu, C., Echols, H., Flanagan, J., Hayer, M.K., and Hartl, F.U. (1992). Successive action of DnaK, DnaJ and GroEL along the pathway of chaperone-mediated protein folding. Nature 356, 683–689.

Levey, T., Westhoff, P., and Meierhoff, K. (2014). Expression of a nuclear-encoded *psbH* gene complements the plastidic RNA processing defect in the PSII mutant *hcf107* in *Arabidopsis thaliana*. Plant J. 80, 292–304.

Li, H., Bai, M., Jiang, X., Shen, R., Wang, H., Wang, H., and Wu, H. (2020). Cytological evidence of BSD2 functioning in both chloroplast division and dimorphic chloroplast formation in maize leaves. BMC Plant Biol. 20, 17.

Liu, C., Young, A.L., Starling-Windhof, A., Bracher, A., Saschenbrecker, S., Rao, B.V., Rao, K.V., Berninghausen, O., Mielke, T., Hartl, F.U., Beckmann, R., and Hayer-Hartl, M. (2010). Coupled chaperone action in folding and assembly of hexadecameric Rubisco. Nature 463, 197–202.

Minai, L., Wostrikoff, K., Wollman, F.A., and Choquet, Y. (2006). Chloroplast biogenesis of photosystem II cores involves a series of assembly-controlled steps that regulate translation. The Plant Cell 18, 159–175.

Mizohata, E., Matsumura, H., Okano, Y., Kumei, M., Takuma, H., Onodera, J., Kato, K., Shibata, N., Inoue, T., Yokota, A., and Kai, Y. (2002). Crystal structure of activated ribulose-1,5-bisphosphate carboxylase/oxygenase from green alga *Chlamydomonas reinhardtii* complexed with 2-carboxyarabinitol-1,5-bisphosphate. J. Mol. Biol. 316, 679–691.

Mühlbauer, S.K., and Eichacker, L.A. (1998). Light-dependent Formation of the Photosynthetic Proton Gradient Regulates Translation Elongation in Chloroplasts. J. Biol. Chem. 273, 20935–20940.

Onishi, M., and Pringle, J.R. (2016). Robust Transgene Expression from Bicistronic mRNA in the Green Alga *Chlamydomonas reinhardtii*. G3 6, 4115–4125.

Onizuka, T., Endo, S., Akiyama, H., Kanai, S., Hirano, M., Yokota, A., Tanaka, S., and Miyasaka, H. (2004). The rbcX gene product promotes the production and assembly of ribulose-1,5-bisphosphate carboxylase/oxygenase of *Synechococcus sp. PCC7002* in *Escherichia coli*. Plant Cell Physiol. 45, 1390–1395.

Rodermel, S., Haley, J., Jiang, C.Z., Tsai, C.H., and Bogorad, L. (1996). A mechanism for intergenomic integration: abundance of ribulose bisphosphate carboxylase small-subunit protein influences the translation of the large-subunit mRNA. Proc. Natl. Acad. Sci. USA 93, 3881–3885.

Rohr, M., Ries, F., Herkt, C., Gotsmann, V.L., Westrich, L.D., Gries, K., Trosch, R., Christmann, J., Chaux-Jukic, F., Jung, M., Zimmer, D., Muhlhaus, T., Sommer, F., Schroda, M., Keller, S., Mohlmann, T., and Willmund, F. (2019). The Role of Plastidic Trigger Factor Serving Protein Biogenesis in Green Algae and Land Plants. Plant Physiol. 179, 1093–1110.

Roth, R., Hall, L.N., Brutnell, T.P., and Langdale, J.A. (1996). bundle sheath defective2, a Mutation That Disrupts the Coordinated Development of Bundle Sheath and Mesophyll Cells in the Maize Leaf. Plant Cell 8, 915–927.

Roy, H., Bloom, M., Milos, P., and Monroe, M. (1982). Studies on the assembly of large subunits of ribulose bisphosphate carboxylase in isolated pea chloroplasts. J. Cell Biol. 94, 20–27.

Salesse-Smith, C.E., Sharwood, R.E., Busch, F.A., Kromdijk, J., Bardal, V., and Stern, D.B. (2018). Overexpression of Rubisco subunits with RAF1 increases Rubisco content in maize. Nat. Plants 4, 802–810.

Salesse, C., Sharwood, R., Sakamoto, W., and Stern, D. (2017). The Rubisco Chaperone BSD2 May Regulate Chloroplast Coverage in Maize Bundle Sheath Cells. Plant Physiol. 175, 1624–1633.

Sambrook, J., Fritsch, E.F., and Maniatis, T. (1989). Molecular cloning: a laboratory manual. (Cold spring harbor laboratory press).

Saschenbrecker, S., Bracher, A., Rao, K.V., Rao, B.V., Hartl, F.U., and Hayer-Hartl, M. (2007). Structure and function of RbcX, an assembly chaperone for hexadecameric Rubisco. Cell 129, 1189–1200.

Shapira, M., Lers, A., Heifetz, P.B., Irihimovitz, V., Barry Osmond, C., Gillham, N.W., and Boynton, J.E. (1997). Differential regulation of chloroplast gene expression in Chlamydomonas reinhardtii during photoacclimation: light stress transiently suppresses synthesis of the Rubisco LSU protein while enhancing synthesis of the PS II D1 protein. Plant Mol. Biol. 33, 1001–1001.

Tabita, F.R., Satagopan, S., Hanson, T.E., Kreel, N.E., and Scott, S.S. (2008). Distinct form I, II, III, and IV Rubisco proteins from the three kingdoms of life provide clues about Rubisco evolution and structure/function relationships. J. Exp. Bot. 59, 1515–1524.

Tarnawski, M., Gubernator, B., Kolesinski, P., and Szczepaniak, A. (2008). Heterologous expression and initial characterization of recombinant RbcX protein from *Thermosynechococcus elongatus* BP-1 and the role of RbcX in RuBisCO assembly. Acta Biochim. Pol. 55, 777–785.

Vitlin Gruber, A., and Feiz, L. (2018). Rubisco Assembly in the Chloroplast. Front Mol. Biosci. 5, 24 (21-11).

Whitney, S.M., Birch, R., Kelso, C., Beck, J.L., and Kapralov, M.V. (2015). Improving recombinant Rubisco biogenesis, plant photosynthesis and growth by coexpressing its ancillary RAF1 chaperone. Proc. Natl. Acad. Sci. USA 112, 3564–3569.

Wittig, I., and Schagger, H. (2005). Advantages and limitations of clear-native PAGE. Proteomics 5, 4338–4346.

Wollman, F.A., Minai, L., and Nechushtai, R. (1999). The biogenesis and assembly of photosynthetic proteins in thylakoid membranes1. Biochim. Biophys. Acta 1411, 21–85.

Woodson, J.D., and Chory, J. (2008). Coordination of gene expression between organellar and nuclear genomes. Nat. Rev. Genet. 9, 383–395.

Wostrikoff, K., and Stern, D. (2007). Rubisco large-subunit translation is autoregulated in response to its assembly state in tobacco chloroplasts. Proc. Natl. Acad. Sci. USA 104, 6466–6471.

Wostrikoff, K., Girard-Bascou, J., Wollman, F.A., and Choquet, Y. (2004). Biogenesis of PSI involves a cascade of translational autoregulation in the chloroplast of Chlamydomonas. EMBO J. 23, 2696–2705.

Wostrikoff, K., Clark, A., Sato, S., Clemente, T., and Stern, D. (2012). Ectopic Expression of Rubisco Subunits in Maize Mesophyll Cells Does Not Overcome Barriers to Cell Type-Specific Accumulation. Plant Phys. 160, 419–432.

Xia, L.Y., Jiang, Y.L., Kong, W.W., Sun, H., Li, W.F., Chen, Y., and Zhou, C.Z. (2020). Molecular basis for the assembly of RuBisCO assisted by the chaperone Raf1. Nat. Plants.

Yosef, I., Irihimovitch, V., Knopf, J.A., Cohen, I., Orr-Dahan, I., Nahum, E., Keasar, C., and Shapira, M. (2004). RNA binding activity of the ribulose-1,5-bisphosphate carboxylase/oxygenase large subunit from Chlamydomonas reinhardtii. J. Biol. Chem. 279, 10148–10156.

Zhan, Y., Dhaliwal, J.S., Adjibade, P., Uniacke, J., Mazroui, R., and Zerges, W. (2015). Localized control of oxidized RNA. J. Cell Sci. 128, 4210–4219.

